# A highly prevalent and pervasive densovirus discovered among sea stars from the North American Atlantic Coast

**DOI:** 10.1101/781609

**Authors:** Elliot W. Jackson, Charles Pepe-Ranney, Mitchell R. Johnson, Daniel L. Distel, Ian Hewson

## Abstract

Viral metagenomes prepared from tissues from Forbes’ sea star (*Asterias forbesi)* led to the discovery of a complete genome of a novel sea star densovirus (AfaDV). The genome organization of AfaDV and phylogenetic analysis place this virus among the *Ambidensovirus* genus in the subfamily *Densoviridae*, family *Parvoviridae*. AfaDV shares 78% nucleotide pairwise identity to the sea star associated densovirus (SSaDV), previously described as the putative causative agent of Sea Star Wasting Syndrome among sea stars from the Northwest Pacific. SSaDV was not found in specimens collected in this study, and the discovery of AfaDV might explain previous reports of SSaDV among sea stars from the Atlantic Coast. A qPCR assay was designed to assess tissue tropism, host specificity, and prevalence of AfaDV among wild populations of sea stars at five locations on the North American Atlantic Coast. AfaDV was detected in all three common sea star species (*Asterias forbesi, Asterias rubens*, and *Henricia sp*.) found in the region and was highly prevalent (80-100% of individuals tested, n=134), among populations collected at disparate sites 7 years apart. AfaDV was detected in the body wall, gonads, and pyloric caeca (digestive gland) of specimens but was not detected in their coelomic fluid. A significant difference in viral load was found between tissue types with the pyloric caeca having the highest viral load suggesting it is the primary site of viral replication in the animal. Further investigation of *Asterias forbesi* gonad tissue found germline cells (oocytes) to be virus positive suggesting a potential route of vertical transmission. Taken together, these observations show that the presence AfaDV is not an indicator of Sea Star Wasting Syndrome because AfaDV is a common constituent of these animals’ microbiome, regardless of health. These results broaden the understanding of echinoderm densoviruses outside the context of disease that suggest these viruses might form commensal or mutualistic relationships with their hosts.

## Introduction

Densoviruses, also known as densonucleosis viruses, are icosahedral, non-enveloped viruses that have monopartite linear single-stranded DNA genomes that are typically 4-6 kb packaged in a 20-25 nm diameter capsid shell (1, 2). Densoviruses are a subfamily (*Densovirinae)* within the greater family *Parvoviridae* and are known to infect arthropods, specifically insects and shrimp (1). Prior to the advent of high throughput sequencing technology, the discovery of densoviruses was driven by investigations of epizootics occurring in laboratory populations and breeding facilities of economically important invertebrates (e.g. silkworms, crickets, shrimp) or through infected cell lines (e.g. mosquito C6/36 cell line) (3–7). The majority of densoviruses isolated to date share a common pathology, causing hypertrophied nuclei in affected tissues, and are generally more virulent at early life stages of its host (1). Densoviruses have also been shown to be mutualists. For example, sublethal infections in rosy apple aphids are correlated with a winged phenotype that has a lower fecundity compared to non-winged aphids (8). The cost of infection lowers the fecundity of the individual to promote the growth of wings that increases mobility and the potential for the host and the virus to disperse (8). Although densoviruses have been primarily studied in insects and crustaceans, analysis of transcriptomic datasets and viral metagenomes prepared from metazoan tissues have found endogenous and exogenous densovirus sequences from a much wider host range (9–13). These finding suggest that densoviruses may be common constituents of many invertebrate viromes.

Densovirus sequences have recently been recovered from echinoderm tissues and have been implicated as potential pathogens though their relationship to echinoderms is unknown (10, 11, 14). In 2013-2015 a mortality event termed sea star wasting disease or syndrome (SSWS) (also referred to as Asteroid Idiopathic Wasting Syndrome) affected >20 sea star species along a broad geographic range from California to Alaska. A densovirus (<u>S</u>ea <u>S</u>tar <u>a</u>ssociated <u>D</u>ensovirus or SSaDV) was discovered and hypothesized to cause SSWS (11, 14). Previous work has reported the presence of SSaDV, using PCR and qPCR (11, 15, 16), among sea stars on the North American Atlantic Coast though was not found to be significantly correlated with SSWS (15, 16). The cause of SSWS among echinoderms on the Atlantic Coast is however hypothesized to be viral in nature (16).

We sought to investigate the presence of SSaDV and more generally survey the viral diversity of densoviruses in sea stars inhabiting the North American Atlantic Coast. Viral metagenomes were prepared from *Asterias forbesi*, a common sea star found in the sub-tidal environment from the North American Atlantic Coast. This led to the discovery of a complete genome of a novel sea star densovirus hereafter referred to as *<u>A</u>sterias forbesi* <u>a</u>ssociated <u>d</u>enso<u>v</u>irus or AfaDV. AfaDV is the second sea star associated densovirus discovered among sea stars thus far. In contrast to previous work, we did not find any evidence of SSaDV among sea stars on the Atlantic Coast through metagenomic analysis and PCR surveys (11, 15, 16). Using qPCR and PCR, we investigated the geographic distribution of AfaDV, the host specificity, tissue tropism, and investigated a potential route of transmission. Our results show that AfaDV has a broad geographic range, is not species specific, has a wide tissue tropism, and is potentially vertically transmitted.

## Methods

### Viral metagenomic preparation and bioinformatic analysis

Viral metagenomes were prepared using a protocol that was adapted and modified from existing laboratory protocols (17). The viral metagenomes used in this study have been previously analyzed and reported for the presence of circular ssDNA viruses (18). Six *Asterias forbesi* that displayed signs characteristic of SSWS (arm detachment, mucoid appearance on aboral surface, disintegration of epidermal tissue) were collected from Canoe Beach, Nahant Bay, Massachusetts, USA (42.420889, −70.906416) in September/October 2015 (19). Animals were flash frozen in liquid nitrogen upon collection then stored at −80°C until dissection. The pyloric caeca and body wall tissue from these animals were pooled separately in 0.02μm-filtered 1X PBS then homogenized in a bleached-cleaned NutriBullet for 60s. Tissue homogenates were pelleted by centrifugation at 3,000 × *g* for 180s, and the supernatant was syringe filtered through Millipore Sterivex-GP 0.22μm polyethersulfone filters into bleach-treated and autoclaved Nalgene Oak Ridge High-Speed Centrifugation Tubes. Filtered homogenates were adjusted to a volume of 35mL by adding 0.02μm-filtered 1X PBS and amended with 10% (wt/vol) PEG-8000 and precipitated for 20 hours at 4°C. Insoluble material was then pelleted by centrifugation at 15,000 × *g* for 30 minutes. The supernatant was decanted and pellets were resuspended in 1mL of 0.02μm-filtered nuclease-free H_2_O. The samples were treated with 0.2 volumes (200μl) of CHCl_3_, inverted three times, and incubated at room temperature for 10 minutes. After a brief centrifugation, 800μl of supernatant was transferred into 1.5mL microcentrifuge tube. Samples were treated with 1.5μl of TURBO DNase (2U/μl) (Invitrogen), 1μl of RNase One (10U/μl) (Thermo Scientific), and 1μl of Benzonase Nuclease (≥250U/μl) (MilliporeSigma) and incubated at 37 °C for 3 hours. 0.2 volumes (160μl) of 100mM EDTA was added to the sample after incubation. Viral DNA was extracted from 500-μl subsamples using the Zymo Research Viral DNA kit following the manufacturer’s protocol and subsequently amplified isothermally at 30°C using Genomiphi Whole Genome Amplification Kit (GE Healthcare, Little Chalfont, UK). Samples were cleaned and concentrated using a Zymo Research DNA Clean and Concentrator kit, and DNA was quantified using Pico Green reagents. Samples were prepared for Illumina sequencing using the Nextera XT DNA Library Preparation Kit (Illumina, San Diego, CA, USA) prior to 2×250bp paired-end Illumina MiSeq sequencing at Cornell University Core Laboratories Center (Ithaca, NY, USA).

Libraries generated from both samples (pyloric caeca and body wall) were first interleaved into one file. Reads were then trimmed for read quality and Illumina adapters and filtered for phiX contamination then merged and normalized to a target depth of 100 and a minimum depth of 1 with an error correction parameter. Read quality filtering, trimming, contamination removal, merging, normalization, and read mapping were done using the BBtools suite (20). The merged and unmerged read normalized libraries were used for *de novo* assembly using SPAdes (21). Contigs less than 500nt were discarded after assembly and the remaining contigs were subject to tBLASTx against a curated in-house database of ssDNA viruses (22). Contigs with significant (*e*-value < 1×10^−8^) sequence similarity to SSaDV (“<u>S</u>ea <u>S</u>tar <u>a</u>ssociated <u>D</u>ensovirus”) were isolated and reads were mapped back to contigs with a minimum identity of 0.95 to obtain average coverage and coverage distribution across the contigs (**Figure 1**). ORFs were defined in Geneious with a minimum size of 550nt, and the hairpin structures in the inverted terminal repeats (ITRs) were determined using Mfold (23, 24). AfaDV genome sequence has been deposited in GenBank under the accession #MN190158. Metagenomic libraries have been deposited in GenBank under BioProject ID PRJNA555067. All statistical and bioinformatics analyses can be found at https://github.com/ewj34/AfaDV-Viral-Metagenome.

**Figure 1:**
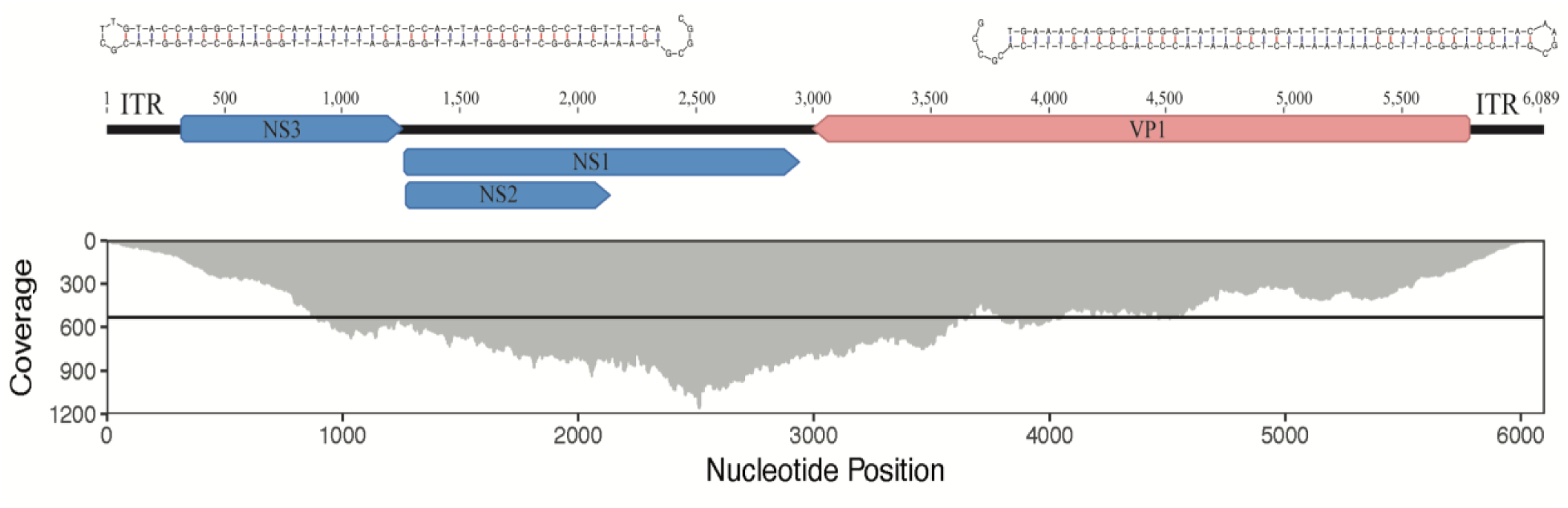
Genome architecture and base coverage of *<u>A</u>sterias forbesi* <u>a</u>ssociated <u>d</u>enso<u>v</u>irus (AfaDV) (top) Structural hairpins 88nt long located in the inverted terminal repeats at the end of the genome. (middle) Genome organization with ORF colored by putative function. Red corresponds to structural protein (VP) and blue corresponds to non-structural proteins (NS1, NS2, NS3). (bottom) Read coverage distribution across genome. Black line indicates 532x average base coverage across genome.

Phylogenetic relationships among AfaDV and 45 densovirus genomes were inferred by a maximum likelihood method, implementing SMS (smart model selection) in PhyML 3.0 using a MUSCLE amino acid sequence alignment of NS1 with default parameters (25, 26). The region of NS1 used for alignment (sequence length 437.7 ± 50; mean ± SD) spanned Motif I of the RC endonuclease domain to Motif C of the SF3 helicase domain (**Supplementary file 3**). Branch support was determined by bootstrapping at 100 iterations. The phylogenetic tree was visualized and edited using iTOL (27).

### Specimen collection and nucleic acid extraction

Three species of sea stars - *Asterias forbesi, Asterias rubens* and *Henricia sp.* - were collected from five locations along the Atlantic Coast of the United States from 2012 to 2019 (**Supplementary Table 1**) (19). 83 *Asterias forbesi*, 16 *Henricia sp*, and 35 *Asterias rubens* were collected totaling 134 sea stars. Prior to vivisection or dissection, animal length was measured by total diameter (i.e. ray to ray). Animals were either vivisected immediately after collection and flash frozen in liquid nitrogen upon collection then stored at −80°C until dissection or cryopreserved and stored at −20°C until dissection. Coelomic fluid was extracted from only animals vivisected using a 25G × 1½ (0.5mm × 25mm) needle attached to a 3 mL syringe inserted through the body wall into the coelomic cavity. Gonads, body wall, and pyloric caeca were collected from all animals vivisected, but not every tissue type was collected from animals that were dissected (**Supplementary Table 1**). In total 368 samples were collected. DNA was extracted from tissues (14 −200mg wet weight) and coelomic fluid (140μl - 1000μl) samples using Zymo Research Tissue & Insect DNA kits or Zymo Research Quick DNA Miniprep Plus Kit following the manufacture’s protocol (**Supplementary Table 1**). All DNA samples used for qPCR were extracted with the Zymo Research Tissue & Insect DNA kit. Coelomocytes in the coelomic fluid were pelleted by centrifugation at 10,000 × *g* for 5 minutes then resuspended in 200μl of 0.02μm-filtered nuclease-free H_2_O prior to DNA extraction. DNA was quantified using Quant-iT PicoGreen dsDNA Assay kit (Invitrogen).

### Genome verification

AfaDV was amplified by PCR with overlapping primers to produce 16 amplicons. Amplicons generated from PCR were gel visualized with ethidium bromide, and the remaining PCR product was cleaned and concentrated using a Zymo Research DNA Clean and Concentrator kit and submitted for Sanger sequencing at Cornell University Core Laboratories Center (Ithaca, NY, USA). The resulting amplicons were assembled to form a contig spanning nucleotide positions 214 to 5860 (92.7% of total genome length). The assembled contig was identical to the contig generated from the *de novo* assembly. This process was repeated on tissue samples from the three sea star species collected to verify the viral genome sequence among each host species. Primers were designed using Primer3 (28). Reaction conditions and primer sequences can be found in **Supplementary Table 2**.

### Cloning

Putative VP and NS genes of AfaDV and SSaDV were inserted into the pGEM-t-Easy vector and used to transform NEB 5-alpha competent cells. Primers, PCR conditions, and plasmid constructs can be found in **Supplemental Table 2/Supplemental Figure 1**. PCR amplicons were gel purified, poly(A) tailed, and ligated to pGEM-t-Easy vectors. The pUC19-SSaDV construct was synthesized by Genscript. Plasmid constructs were verified by Sanger sequencing.

### AfaDV prevalence and viral load

Viral prevalence and viral load were determined using quantitative(q)PCR using TaqMan chemistry. qPCR primers and probe were designed using Primer3 targeting the VP region positions 5634 to 5728 (28). Reaction conditions and primer/probe/oligonucleotide standard sequences can be found in **Supplementary Table 2**. All qPCR reactions, including no-template controls, were performed in duplicate on a StepOnePlus™ Real-Time PCR system (Applied Biosystems). Duplicate eight-fold standard dilutions were included in all qPCR runs. 25μl reaction volumes contained 0.02μl (200 pmol) of each primer and probe, 12.5μl of 2X SsoAdvanced ™ Universal Probes Supermix (Bio-Rad), 10.44μl of nuclease-free H_2_O, and 2μl of template. Viral quantity was calculated by StepOnePlus™ software (version 2.3 Applied Biosystems) by averaging the cycle threshold between duplicates and interpolating values against the standard curve. Viral quantities were adjusted by extraction volume and standardized by sample weight. Samples were considered positive when technical replicates were both positive and Ct standard deviation was <1.0. The lower limit of detection used in this study was 30 copies per reaction (average Ct value of 34) or 40 copies·mg^−1^ tissue. Viral prevalence was defined as the total number of positive samples among a sample type, and viral load was defined as the mean copy number of technical replicates in a positive sample.

### Oocyte collection, DNA extraction, PCR, SSaDV survey

Fifteen *Asterias forbesi* were collected from Woods Hole, Massachusetts in June/July 2019. Gonadal tissue and pyloric caeca were taken through a small incision to the arm and manually extracted with forceps (**Supplemental Table 1**). Isolation of oocytes was performed according to Wessel *et al.*, 2010 (29). Oocytes were isolated by mincing gonadal tissue in 0.2μm filtered sea water then poured through cheesecloth, pelleted by centrifugation, decanted, and pipetted for DNA extraction. DNA extractions were performed using Zymo Research Quick DNA MiniPrep kit Plus. PCR reactions included a kit negative control (i.e. extraction blank) and PCR reagent negative control to account for false-positives. PCR cycle conditions can be found in **Supplemental Table 2.**

Thirty pyloric caeca samples were screened for SSaDV to determine the presence of SSaDV among North Atlantic sea stars. Ten pyloric caeca samples from Woods Hole, Shoals Marine Lab, and Nahant were chosen. The specificity of primers was validated (i.e. no cross amplification observed) by screening against the appropriate plasmid constructs prior to screening DNA extracts (**Supplemental Figure 1**). PCR cycle conditions can be found in **Supplemental Table 2**.

## Results

### Genome analysis and phylogeny

High-throughput sequencing of metaviromes prepared from the pyloric caeca and body wall of sea star samples generated 2.42 × 10^7^ and 7.66 × 10^6^ reads/library respectively totaling 3.18 ×10^7^ reads. SPAdes assembly of trimmed, decontaminated, merged and unmerged, normalized reads resulted in one contig that was 6,089 nt that was significantly (*e*-value < 1×10^−8^) similar to SSaDV (**Figure 1**). Read mapping to this contig recruited 16,737 reads that gave an average base coverage of 532 that proportionately made up 0.052% of the total reads (**Figure 1**). A total of 16,732 reads mapped came from the metavirome prepared from the pyloric caeca and 5 reads mapped from the metavirome prepared from body wall tissue. The contig contained 4 ORFs that putatively encode non-structural proteins (NS1, NS2, and NS3) and a structural protein (VP) (**Figure 1**). As a whole, the contig encodes all components of the NS and VP cassettes that are characteristic of the genus *Ambidensovirus* within the subfamily *Densovirinae.* Phylogenetic analyses (Maximum Likelihood) of the aforementioned sequences indicate that this novel densovirus falls within a well-supported clade that includes other *Ambidensoviruses* and shares a most recent common ancestry with the previously described sea star densovirus, SSaDV (**Figure 2**). AfaDV and SSaDV shared a 77.9% pairwise nucleotide identity across entire genomes though the putative NS1, NS2, and NS3 genes have pairwise nucleotide identities of 88.7%, 88.1% and 74.4% respectively while the putative VP gene share a 77.1% pairwise nucleotide identity. It is highly likely that the contig is a complete genome because of the presence of the hairpins located at the ends of the inverted terminal repeats (ITRs) which are characteristic of parvoviruses (30). The hairpins are 93nt long that form stem loop structures that are thermodynamically favorable (ΔG = −52.71) (**Figure 1**).

**Figure 2:**
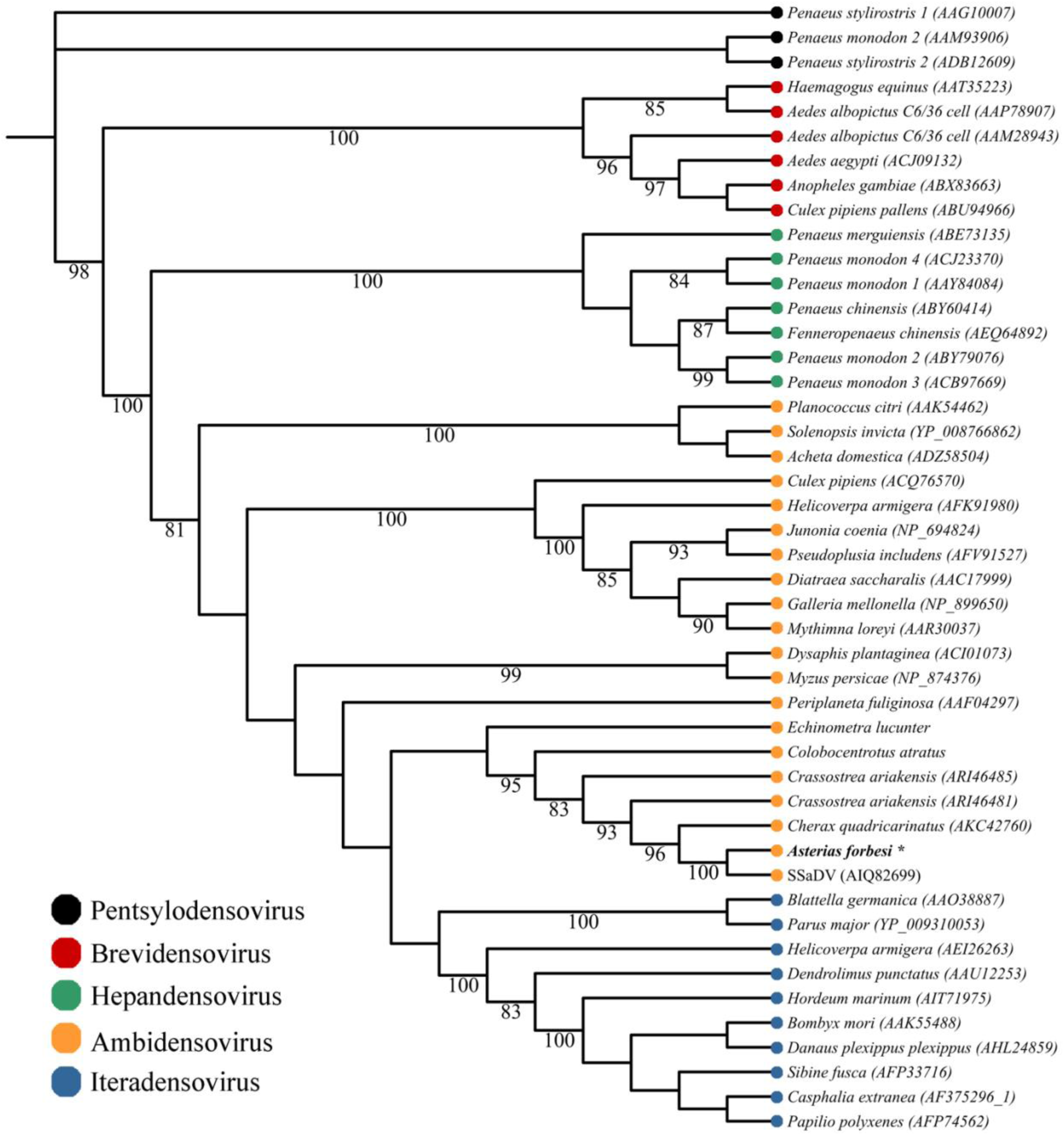
Maximum likelihood phylogeny of densoviruses (AIC; LG +G+I+F). Cladogram is based on an amino acid alignment performed by MUSCLE of the NS1 region spanning Motif I of the RC endonuclease domain to Motif C of the SF3 helicase domain (amino acid sequence length 437.7 ± 50; mean ± SD). Branch support bootstrapped at 100 iterations and terminal node colors correspond to densovirus genus. Italicized names correspond to animal genus and species for which the densovirus was isolated. AfaDV denoted in bold with *.

### Tissue tropism, prevalence, and biogeography

SSaDV was not detected in any of the pyloric caeca samples screened. AfaDV was detected from sea stars collected from 2012 – 2019 from four of the five locations (Appledore Island, Maine, Nahant, Massachusetts, Woods Hole, Massachusetts, and the Mystic Aquarium, Connecticut)(**Figure 3**). AfaDV was not detected from 6 *Asterias forbesi* collected from Bar Harbor, ME (**Figure 3**). The overlapping PCR primer set successfully amplified AfaDV sequences from the three sea star species collected in this study. To assess potential tissue tropism, we screened pyloric caeca, gonads, body wall and coelomic fluid from animals via qPCR. AfaDV was detected more frequently in the pyloric caeca (86%) > body wall (70%) > gonads (57%) and was not detected in any coelomic fluid samples (**Figure 4**). Viral load was significantly different among tissue types (Welch’s ANOVA, F_2,117.66_ = 29.205, *p* = 5.028 × 10^−11^; **Figure 4**). AfaDV log_10_ transformed values was significantly greater in the pyloric caeca (3.3 ± 0.88; mean ± SD) than in the body wall (2.6 ± 0.44, Games-Howell, *p* = 5.6 × 10^−8^) and gonads (2.3 ± 0.28, Games-Howell, *p* = 5.2 × 10^−11^). AfaDV load was not significantly different between gonads and body wall (Games-Howell, *p* = 0.13). Viral load was also significantly correlated with animal length (Linear regression, *p* = 1.26 × 10^−7^) though animal length had a low degree of correlation (R_2_ = 0.142) (**Figure 4**). Average DNA concentration differences found between tissue types did not reflect the trends found in viral loads across tissue types (**Supplemental Figure 2**).

**Figure 3:**
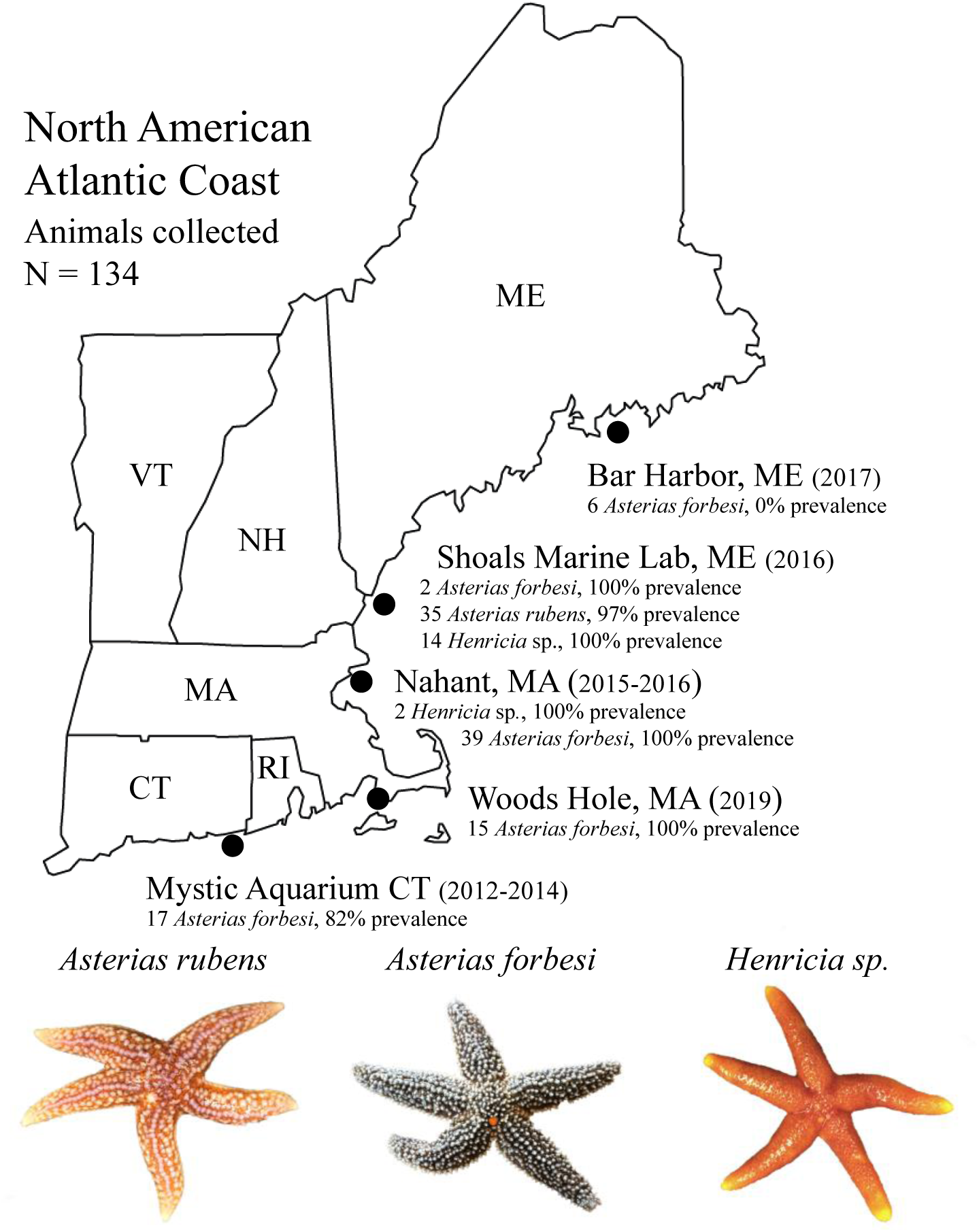
AfaDV prevalence among sea star populations along the North American Atlantic Coast. A total of 134 animals from three species of sea stars were screened via qPCR or PCR for AfaDV. Year of sampling shown in parentheses. Prevalence for each species corresponds to the number of animals positive for AfaDV divided by the total listed next to each species.

**Figure 4:**
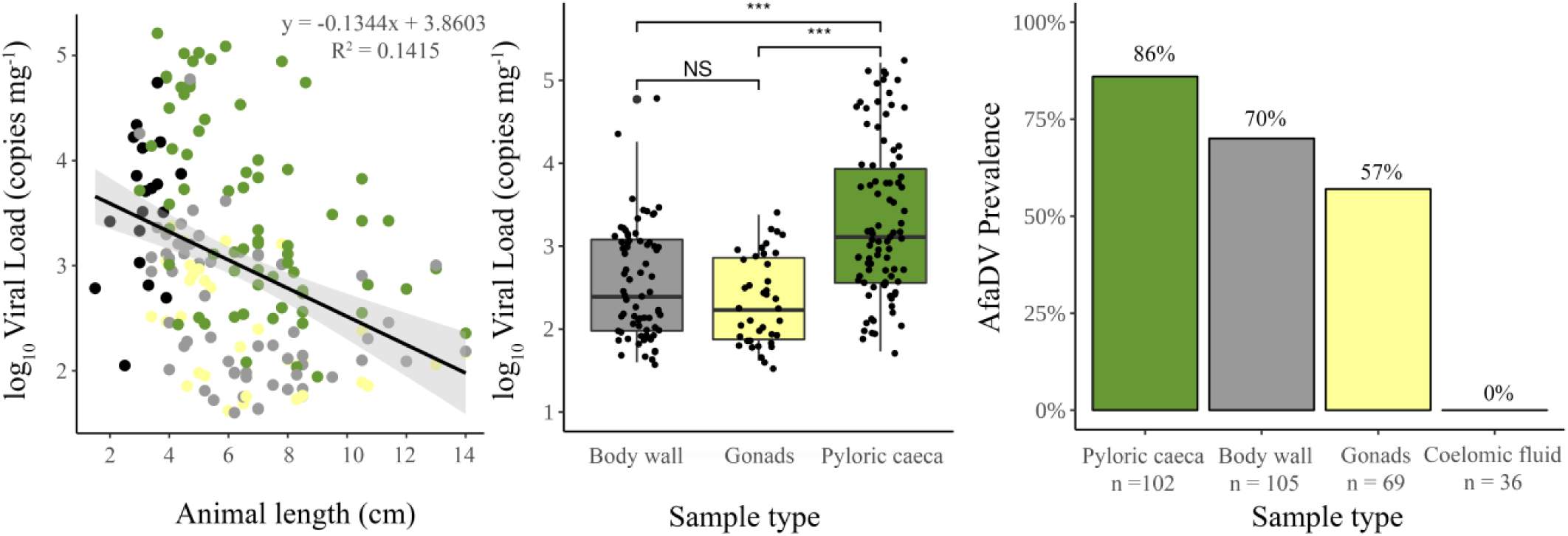
Viral load and tissue prevalence of AfaDV (left) Linear regression analysis of viral load and animal length reported as total diameter. Colors correspond to sample type. Black dots represent cross-section samples; (middle) Viral load comparison between tissue type. (right); Prevalence of AfaDV among tissue types. *** *p* = < 0.001. NS = No significance.

### Vertical transmission of AfaDV

The pyloric caeca of 15 *Asterias forebsii* from Woods Hole, Massachusetts were virus positive via PCR. 5/10 oocytes isolated from females were virus positive and 2/5 gonadal tissue from male sea stars were virus positive (**Figure 5, Supplemental Figure 3**).

**Figure 5:**
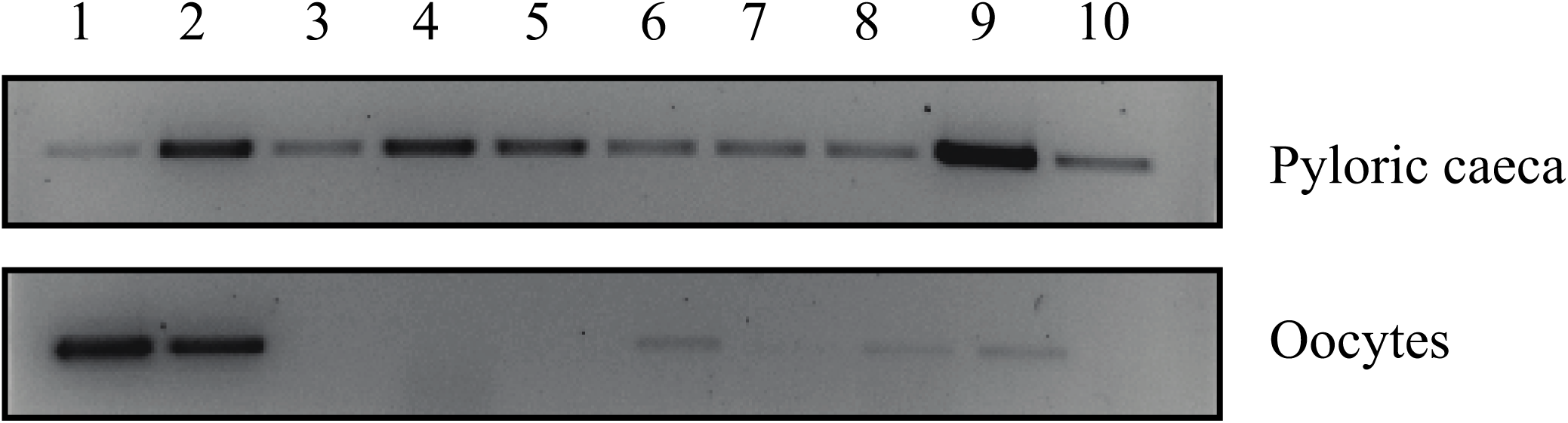
PCR detection of AfaDV from DNA extracted from pyloric caeca and oocytes collected from 10 female *Asterias forbesi* from Woods Hole, Massachusetts.

## Discussion

The discovery of SSaDV and detection in sea stars from the Northwest Pacific and Northwest Atlantic suggest that SSaDV is associated with SSWS in disparate geographic regions (11). Subsequent investigations (15, 16) have supported an association between the occurrence of SSaDV and the incidence of SSWS among sea stars in the Atlantic using primers that were presumably specific to SSaDV. Previous documentation of SSaDV in Atlantic sea stars however may be confounded by spurious amplification and/or the presence of a genetically similar densovirus. Indeed, the nucleotide similarity of AfaDV and SSaDV suggests that previous primers may have been insufficient in distinguishing these two genotypes which led to the conclusion that SSaDV is associated with sea stars on the Atlantic Coast. By validating the specificity of our primers, we tested the presence of both genotypes and did not find evidence for the presence of SSaDV in sea stars on the Atlantic Coast. These results suggest that SSaDV is limited to sea stars in Northwest Pacific which implies any correlation to SSWS outside of this region, and the sea stars that are found in it, is unlikely. Although SSWS is broadly defined and is most notably observed in the Northwest Pacific, it has been observed elsewhere (31). If densoviruses are correlated with SSWS in disparate geographic regions, unique densovirus genotype(s) may also exist in those regions. Further efforts to document the diversity and biogeography of these viruses may help to elucidate these correlations.

Currently, it is unclear what environmental or host-specific factors shape the biogeography of AfaDV and SSaDV, but the extent of these viruses among wild populations may be underappreciated. Similar to SSaDV, AfaDV is not associated with one species and can be found across a large geographic range (**Figure 3**). To accurately document the prevalence of AfaDV, we first investigated tissue tropism, which has not been established for this virus-host system, to establish the best tissue type for viral detection (Figure 4). Densoviruses typically have a wide tissue tropism in arthropods and can actively replicate in most tissues though replication can be exclusively limited to certain tissues depending on the viral genotype. For example, densoviruses in the genus *Iteradensovirus* replicate exclusively and/or predominantly in midgut epithelium cells of their hosts while *G. mellonella* and *J.coenia* ambidensoviruses replicate in almost all tissues except the midgut epithelium (1, 32). Using qPCR and PCR, we detected AfaDV in the pyloric caeca, gonads, and body wall with varying degrees of prevalence but no positive detection was found in the coelomic fluid (**Figure 4**). It should be noted that our investigation of tissue tropism did not include stomach, intestinal caeca, and radial nerves. Further analysis of these tissues would be needed to establish complete tissue tropism in addition to testing for viral replication among these tissues. Comparative analysis across tissue types showed significantly higher viral loads per unit sample weight in the pyloric caeca than other tissue types (**Figure 4**). This difference was not found to be a reflection in DNA concentrations between tissue types which suggests the viral loads across tissue types is due to a biological difference (**Supplemental Figure 2**). It is not clear whether viral replication is also significantly greater in the pyloric caeca but such differences might result from the susceptibility of cell types, rates of cellular division among tissues or accessibility of tissue to the host immune system. Given that densoviruses replicate during the S-1 phase of mitosis, the trends in viral load across tissues likely reflect differences in cellular proliferation between these tissues (30). We hypothesize that the pyloric caeca has a larger proportion of dividing cells relative to other tissues, thereby explaining the observed differences in viral load. Similarly, the correlation between viral load and animal length could be a reflection of a greater proportion of cellular division in growing individuals.

The detection of AfaDV in the same populations over a two to three-year time span suggest that sea stars can maintain persistent densovirus infections. Persistent infections are common among vertebrate parvoviruses and densoviruses (33–36). Parvoviruses replicate passively and are generally more virulent in fetal and juvenile organisms while adults can maintain persistent infections without showing any clinical signs. For example, shrimp densoviruses can cause acute infections in juveniles but individuals that survive can carry the virus for life transmitting it vertically and horizontally (37). Infected adults rarely show signs of disease even in individuals with heavy infections (36). Persistent infections have been reported in other virus-host invertebrate systems. Notably viruses that infect *Apis mellifera*, the European honey bee, form persistent infections that can lead to acute infections under certain conditions (38, 39). The persistence of AfaDV among host populations brings into question the capacity of sea star densoviruses to cause acute infections under particular circumstances. Such circumstances might include one or a combination of biotic and abiotic factors such as the nutritional status of the host, temperature fluctuations, microbial dysbiosis, or the presence of a vector (e.g. *Varroa destructans*). Considering the high prevalence of AfaDV or SSaDV among sea stars with no obvious signs of infection, the association to SSWS, if any, may be the result of a combined effect of viral infection(s) and biotic or abiotic changes.

The presence of AfaDV in DNA extracted from sea star oocytes suggest that viruses can infect the animal’s germline cells (**Figure 5**). Active or latent infection during embryonic development, larval growth, and metamorphosis could facilitate vertical transmission, an efficient mechanism to achieve high prevalence in adults. Further experimental and microscopic evidence will be necessary to fully establish this route of transmission. The observation of AfaDV in oocytes also suggests these cells may be permissive. Because the whole genome of AfaDV was recovered *in silico*, a synthetic clone could be constructed and microinjected into developing embryos to test the permissiveness of these cells. Given that echinoderms are model organisms in developmental biology and infectious clones are routinely used to study densovirus biology, the tractability of this approach is promising (40–44, 29). These experiments are currently being tested using a SSaDV clone as part of an ongoing investigation to further understand the relationship of densoviruses to sea stars.

Here we report the discovery of a novel sea star densovirus from *Asterias forbesi.* Phylogenetic analysis demonstrates that this virus is closely related to the previously discovered sea star densovirus, SSaDV. Our investigation did not find evidence for the presence of SSaDV in specimens from the North Atlantic, suggesting that SSaDV is limited to Northwest Pacific sea stars. PCR based approaches were used to investigate tissue tropism, prevalence among healthy sea star populations in the environment, and potential vertical transmission of AfaDV. We found AfaDV to have a broad geographic range that spreads across Connecticut to Maine and is found in high prevalence among populations. The vertical transmission of AfaDV may explain these high prevalence rates. The results of this study further our understanding of the association between densoviruses and echinoderms beyond the context of disease. The prevalence and pervasiveness of AfaDV among wild populations suggests that these viruses might form commensal or mutualistic relationships with their hosts. The pathogenicity and interaction of AfaDV at the cellular, larval and adult stage cannot be inferred from these data alone, but it appears that densoviruses may be common constituents of these animal’s microbiome.

## Acknowledgments

The authors thank Dr. Holly Lutz, Taylor Dueweke, Dr. Lisa Abbo, Dr. Charlotte Seid, Dr. Kathy Tuxbury, Kathryn Bland, and Dr. James Coyer for their assistance in animal collection. The staff at Ocean Genome Legacy, Shoals Marine Laboratory, and the Marine Biological Laboratory at Woods Hole. The authors also thank Dr. Kalia Bistolas, Dr. Gary Wessel, and Dr. Nathalie Oulhen for their assistance and suggestions with laboratory work and manuscript comments.

