## Supplemental Figure 1 for "A highly prevalent and pervasive densovirus discovered among sea stars from the North American Atlantic Coast"

**Supplemental Figure 1** – Plasmid constructs created for primer specificity analysis. A) Densovirus genome architecture. ORFs colored and labeled by putative function. Red represents structural proteins (VP) and blue represents non-structural proteins (NS) B) Plasmid constructs created using pGEM-t-Easy.

A)

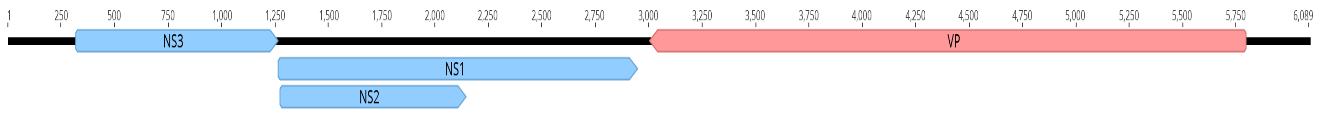


B)

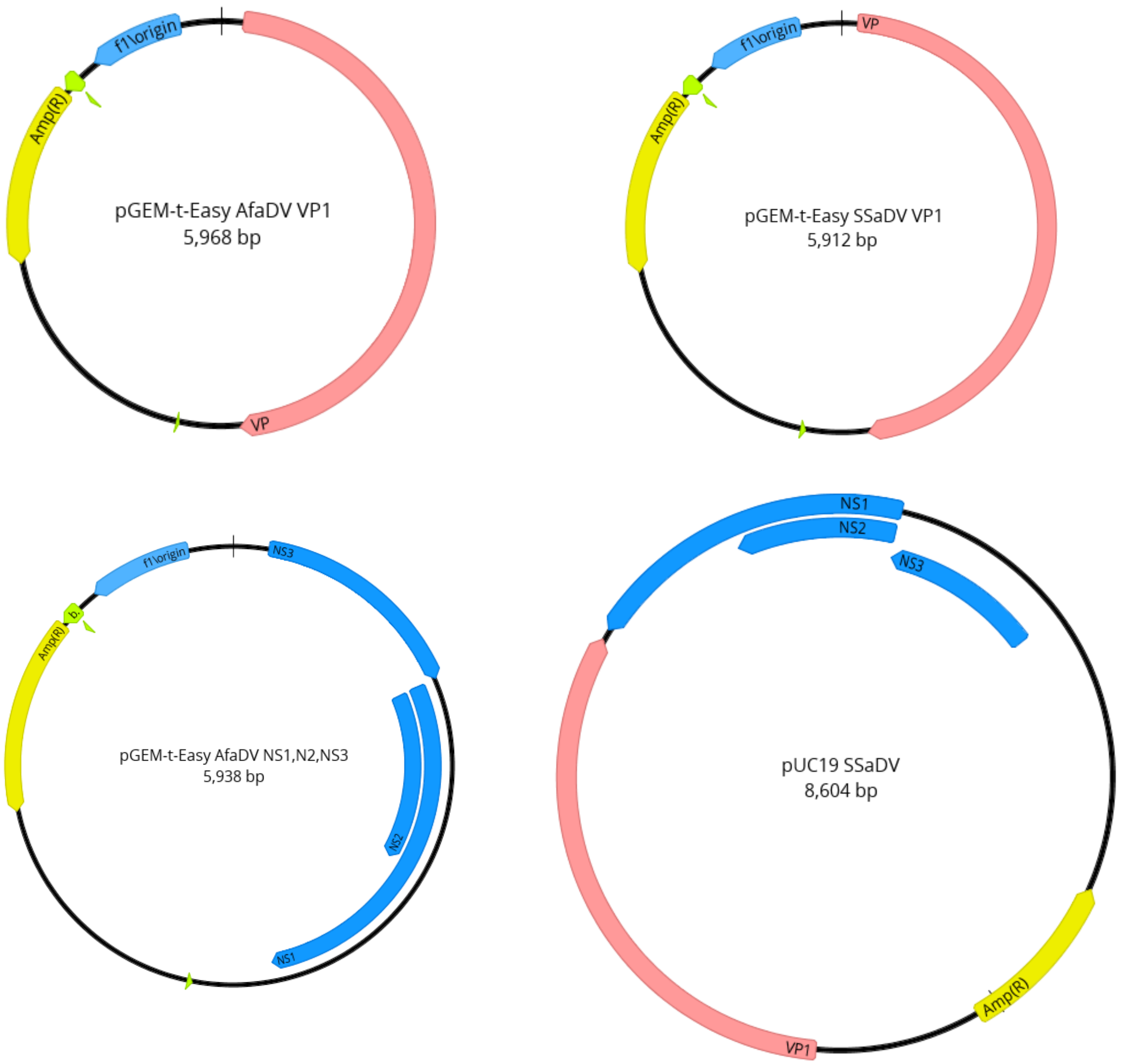
