## Supplemental Figure 2 for "A highly prevalent and pervasive densovirus discovered among sea stars from the North American Atlantic Coast"

**Supplemental Figure 2:** Mean DNA concentration across samples types. Error bars represent two standard errors from the mean.

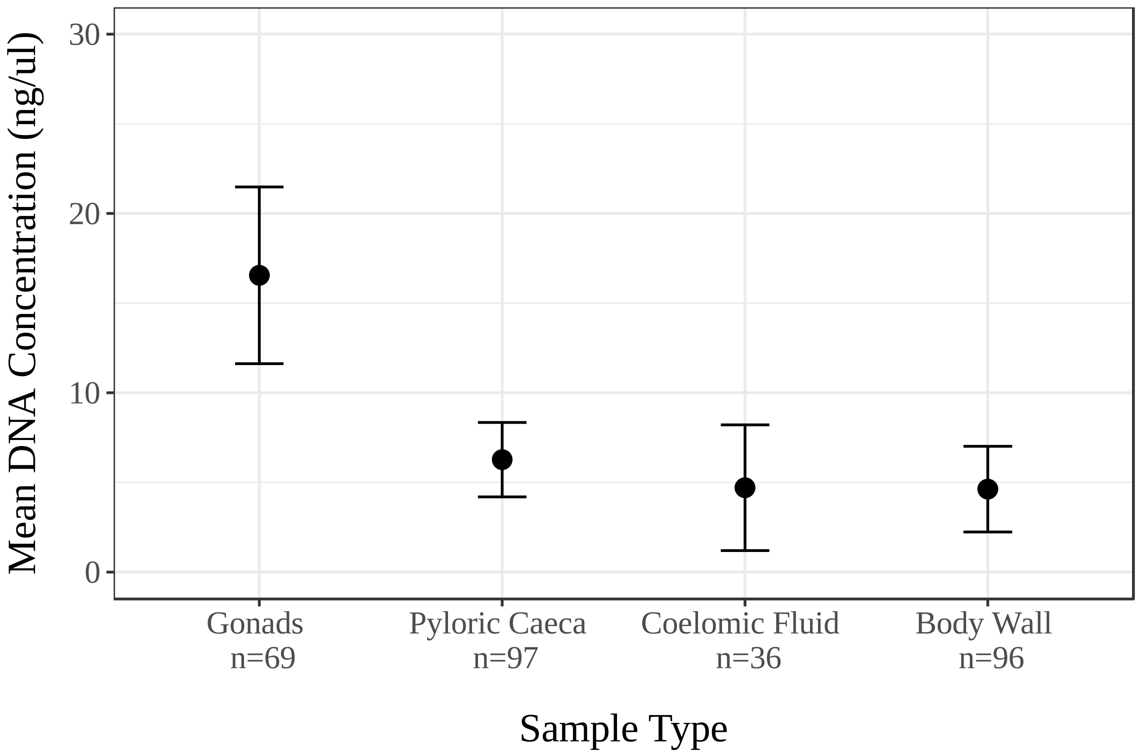
