## Supplemental Figure 3 for "A highly prevalent and pervasive densovirus discovered among sea stars from the North American Atlantic Coast"

**Supplemental Figure 3:** Extended Figure 5. PCR detection of AfDV from DNA extracted from pyloric caeca and oocytes collected from 10 female *Asterias forbesi* from Woods Hole, Massachusetts. B = kit extraction blank for non-template control PCR.

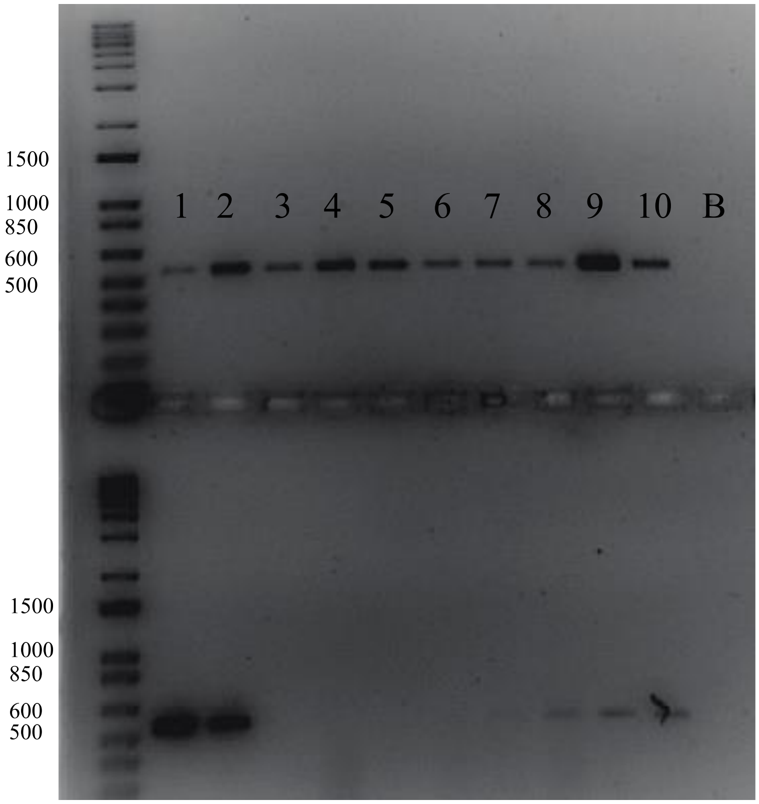
