## Supplemental Table 1 for "A highly prevalent and pervasive densovirus discovered among sea stars from the North American Atlantic Coast"

| Sample# | Animal# | CollectionLocation | CollectionYear | Species | SampleType | Health | Sex | AnimalSize | CollVessel | OGLSampleID | Subsample(mg) | DNAExtractionKit | DNAconc(ng/ul) | Analysis | LogViralLoad |
|---|---|---|---|---|---|---|---|---|---|---|---|---|---|---|---|
| OGL1 | 1 | Nahant, MA | July 7, 2016 | Asterias forbesi | Pyloric caeca | Healthy | NA | 6 | 2 mL | S23233 | 64.80 | Zymo Tissue & Insect | 12.33 | qPCR | 3.71 |
| OGL2 | 1 | Nahant, MA | July 7, 2016 | Asterias forbesi | Gonads | Healthy | NA | 6 | 2 mL | S23233 | 77.60 | Zymo Tissue & Insect | 49.64 | qPCR | 1.62 |
| OGL3 | 1 | Nahant, MA | July 7, 2016 | Asterias forbesi | Body Wall | Healthy | NA | 6 | 2 mL | S23233 | 70.30 | Zymo Tissue & Insect | 1.02 | qPCR | 2.09 |
| OGL4 | 2 | Nahant, MA | July 7, 2016 | Asterias forbesi | Pyloric caeca | Healthy | NA | 5 | 2 mL | S23234 | 64.60 | Zymo Tissue & Insect | 1.36 | qPCR | 4.28 |
| OGL5 | 2 | Nahant, MA | July 7, 2016 | Asterias forbesi | Gonads | Healthy | NA | 5 | 2 mL | S23234 | 78.30 | Zymo Tissue & Insect | 0.56 | qPCR | 1.98 |
| OGL6 | 2 | Nahant, MA | July 7, 2016 | Asterias forbesi | Body Wall | Healthy | NA | 5 | 2 mL | S23234 | 62.40 | Zymo Tissue & Insect | 0.55 | qPCR | 3.02 |
| OGL7 | 3 | Nahant, MA | July 7, 2016 | Asterias forbesi | Pyloric caeca | Healthy | NA | 4.7 | 2 mL | S23235 | 24.90 | Zymo Tissue & Insect | 2.61 | qPCR | 4.75 |
| OGL8 | 3 | Nahant, MA | July 7, 2016 | Asterias forbesi | Gonads | Healthy | NA | 4.7 | 2 mL | S23235 | 74.30 | Zymo Tissue & Insect | 44.59 | qPCR | 3.00 |
| OGL9 | 3 | Nahant, MA | July 7, 2016 | Asterias forbesi | Body Wall | Healthy | NA | 4.7 | 2 mL | S23235 | 75.00 | Zymo Tissue & Insect | 0.24 | qPCR | 4.77 |
| OGL10 | 4 | Nahant, MA | July 7, 2016 | Asterias forbesi | Pyloric caeca | Healthy | NA | 4.8 | 2 mL | S23236 | 75.80 | Zymo Tissue & Insect | 1.60 | qPCR | 4.94 |
| OGL11 | 4 | Nahant, MA | July 7, 2016 | Asterias forbesi | Gonads | Healthy | NA | 4.8 | 2 mL | S23236 | 67.90 | Zymo Tissue & Insect | 0.59 | qPCR | 2.91 |
| OGL12 | 4 | Nahant, MA | July 7, 2016 | Asterias forbesi | Body Wall | Healthy | NA | 4.8 | 2 mL | S23236 | 70.10 | Zymo Tissue & Insect | 0.68 | qPCR | 3.53 |
| OGL13 | 5 | Nahant, MA | July 7, 2016 | Asterias forbesi | Pyloric caeca | Healthy | NA | 4.6 | 2 mL | S23237 | 72.30 | Zymo Tissue & Insect | 5.24 | qPCR | 4.06 |
| OGL14 | 5 | Nahant, MA | July 7, 2016 | Asterias forbesi | Gonads | Healthy | NA | 4.6 | 2 mL | S23237 | 75.00 | Zymo Tissue & Insect | 47.28 | qPCR | 1.86 |
| OGL15 | 5 | Nahant, MA | July 7, 2016 | Asterias forbesi | Body Wall | Healthy | NA | 4.6 | 2 mL | S23237 | 73.20 | Zymo Tissue & Insect | 0.54 | qPCR | 2.28 |
| OGL16 | 6 | Nahant, MA | July 7, 2016 | Henricia sp. | Whole animal | Healthy | NA | 3.3 | 2 mL | S23238 | 66.80 | Zymo Tissue & Insect | 1.30 | qPCR, PCR | 2.81 |
| OGL17 | 7 | Nahant, MA | July 7, 2016 | Henricia sp. | Whole animal | Healthy | NA | 3.9 | 2 mL | S23239 | 74.40 | Zymo Tissue & Insect | 0.59 | qPCR, PCR | 2.69 |
| OGL18 | 8 | Nahant, MA | July 12, 2016 | Asterias forbesi | Whole animal | Healthy | NA | 1.5 | 2 mL | S23240 | 67.20 | Zymo Tissue & Insect | 2.02 | qPCR | 2.79 |
| OGL19 | 9 | Nahant, MA | July 12, 2016 | Asterias forbesi | Whole animal | Healthy | NA | 3 | 2 mL | S23241 | 80.00 | Zymo Tissue & Insect | 0.69 | qPCR | 3.03 |
| OGL20 | 10 | Nahant, MA | July 12, 2016 | Asterias forbesi | Whole animal | Healthy | NA | 2.8 | 2 mL | S23242 | 70.90 | Zymo Tissue & Insect | 1.30 | qPCR | 4.22 |
| OGL21 | 11 | Nahant, MA | July 12, 2016 | Asterias forbesi | Whole animal | Healthy | NA | 3.2 | 2 mL | S23243 | 69.20 | Zymo Tissue & Insect | 3.42 | qPCR | 3.71 |
| OGL22 | 12 | Nahant, MA | July 12, 2016 | Asterias forbesi | Whole animal | Healthy | NA | 2.9 | 2 mL | S23244 | 62.80 | Zymo Tissue & Insect | 2.03 | qPCR | 3.86 |
| OGL23 | 13 | Nahant, MA | July 12, 2016 | Asterias forbesi | Whole animal | Healthy | NA | 3.8 | 2 mL | S23245 | 69.30 | Zymo Tissue & Insect | 0.50 | qPCR | 3.51 |
| OGL24 | 14 | Nahant, MA | July 12, 2016 | Asterias forbesi | Whole animal | Healthy | NA | 4.4 | 2 mL | S23246 | 60.10 | Zymo Tissue & Insect | 0.60 | qPCR | 3.87 |
| OGL25 | 15 | Nahant, MA | July 12, 2016 | Asterias forbesi | Pyloric caeca | Healthy | NA | 3.9 | 2 mL | S23247 | 40.40 | Zymo Tissue & Insect | 1.56 | qPCR | 4.79 |
| OGL26 | 15 | Nahant, MA | July 12, 2016 | Asterias forbesi | Gonads | Healthy | NA | 3.9 | 2 mL | S23247 | 62.20 | Zymo Tissue & Insect | 0.58 | qPCR | 2.47 |
| OGL27 | 15 | Nahant, MA | July 12, 2016 | Asterias forbesi | Body Wall | Healthy | NA | 3.9 | 2 mL | S23247 | 71.10 | Zymo Tissue & Insect | 0.67 | qPCR | 3.11 |
| OGL28 | 16 | Nahant, MA | July 12, 2016 | Asterias forbesi | Pyloric caeca | Healthy | NA | 3.9 | 2 mL | S23248 | 35.50 | Zymo Tissue & Insect | 0.49 | qPCR | 4.80 |
| OGL29 | 16 | Nahant, MA | July 12, 2016 | Asterias forbesi | Gonads | Healthy | NA | 3.9 | 2 mL | S23248 | 77.10 | Zymo Tissue & Insect | 0.69 | qPCR | 3.05 |
| OGL30 | 16 | Nahant, MA | July 12, 2016 | Asterias forbesi | Body Wall | Healthy | NA | 3.9 | 2 mL | S23248 | 62.10 | Zymo Tissue & Insect | 0.50 | qPCR | 3.30 |
| OGL31 | 17 | Nahant, MA | July 12, 2016 | Asterias forbesi | Pyloric caeca | Healthy | NA | 4.1 | 2 mL | S23249 | 62.00 | Zymo Tissue & Insect | 0.59 | qPCR | 4.11 |
| OGL32 | 17 | Nahant, MA | July 12, 2016 | Asterias forbesi | Gonads | Healthy | NA | 4.1 | 2 mL | S23249 | 21.10 | Zymo Tissue & Insect | 0.56 | qPCR | 3.38 |
| OGL33 | 17 | Nahant, MA | July 12, 2016 | Asterias forbesi | Body Wall | Healthy | NA | 4.1 | 2 mL | S23249 | 71.50 | Zymo Tissue & Insect | 0.44 | qPCR | 2.95 |
| OGL34 | 18 | Nahant, MA | July 12, 2016 | Asterias forbesi | Pyloric caeca | Healthy | NA | 4.7 | 2 mL | S23250 | 33.70 | Zymo Tissue & Insect | 0.83 | qPCR | 4.70 |
| OGL35 | 18 | Nahant, MA | July 12, 2016 | Asterias forbesi | Gonads | Healthy | NA | 4.7 | 2 mL | S23250 | 68.10 | Zymo Tissue & Insect | 55.03 | qPCR | 2.86 |
| OGL36 | 18 | Nahant, MA | July 12, 2016 | Asterias forbesi | Body Wall | Healthy | NA | 4.7 | 2 mL | S23250 | 67.20 | Zymo Tissue & Insect | 0.52 | qPCR | 3.21 |
| OGL37 | 19 | Nahant, MA | July 12, 2016 | Asterias forbesi | Pyloric caeca | Healthy | NA | 4.4 | 2 mL | S23251 | 70.20 | Zymo Tissue & Insect | 1.06 | qPCR | 4.70 |
| OGL38 | 19 | Nahant, MA | July 12, 2016 | Asterias forbesi | Gonads | Healthy | NA | 4.4 | 2 mL | S23251 | 76.00 | Zymo Tissue & Insect | 49.21 | qPCR | 2.51 |
| OGL39 | 19 | Nahant, MA | July 12, 2016 | Asterias forbesi | Body Wall | Healthy | NA | 4.4 | 2 mL | S23251 | 67.60 | Zymo Tissue & Insect | 0.59 | qPCR | 3.40 |
| OGL40 | 20 | Nahant, MA | July 12, 2016 | Asterias forbesi | Pyloric caeca | Healthy | NA | 5.4 | 2 mL | S23252 | 66.70 | Zymo Tissue & Insect | 0.96 | qPCR | 4.96 |
| OGL41 | 20 | Nahant, MA | July 12, 2016 | Asterias forbesi | Gonads | Healthy | NA | 5.4 | 2 mL | S23252 | 65.40 | Zymo Tissue & Insect | 0.70 | qPCR | 2.79 |
| OGL42 | 20 | Nahant, MA | July 12, 2016 | Asterias forbesi | Body Wall | Healthy | NA | 5.4 | 2 mL | S23252 | 79.70 | Zymo Tissue & Insect | 0.46 | qPCR | 3.04 |
| OGL43 | 21 | Nahant, MA | July 12, 2016 | Asterias forbesi | Pyloric caeca | Healthy | NA | 5.9 | 2 mL | S23253 | 63.60 | Zymo Tissue & Insect | 1.54 | qPCR | 5.09 |
| OGL44 | 21 | Nahant, MA | July 12, 2016 | Asterias forbesi | Gonads | Healthy | NA | 5.9 | 2 mL | S23253 | 79.60 | Zymo Tissue & Insect | 1.01 | qPCR | 3.23 |
| OGL45 | 21 | Nahant, MA | July 12, 2016 | Asterias forbesi | Body Wall | Healthy | NA | 5.9 | 2 mL | S23253 | 60.20 | Zymo Tissue & Insect | 0.62 | qPCR | 3.62 |
| OGL46 | 22 | Nahant, MA | July 12, 2016 | Asterias forbesi | Pyloric caeca | Healthy | NA | 6.4 | 2 mL | S23254 | 52.50 | Zymo Tissue & Insect | 3.61 | qPCR | 4.53 |
| OGL47 | 22 | Nahant, MA | July 12, 2016 | Asterias forbesi | Gonads | Healthy | NA | 6.4 | 2 mL | S23254 | 73.70 | Zymo Tissue & Insect | 0.54 | qPCR | 2.23 |
| OGL48 | 22 | Nahant, MA | July 12, 2016 | Asterias forbesi | Body Wall | Healthy | NA | 6.4 | 2 mL | S23254 | 71.60 | Zymo Tissue & Insect | 0.46 | qPCR | 3.13 |
| OGL49 | 23 | Nahant, MA | July 12, 2016 | Asterias forbesi | Pyloric caeca | Healthy | NA | 8.6 | 2 mL | S23255 | 66.10 | Zymo Tissue & Insect | 1.42 | qPCR | 4.74 |

| | | | | | | | | | | | | | | | |
|---|---|---|---|---|---|---|---|---|---|---|---|---|---|---|---|
| OGL52 | 24 | Nahant, MA | July 12, 2016 | Asterias forbesi | Whole animal | Healthy | NA | 3.6 | 2 mL | S23256 | 65.30 | Zymo Tissue & Insect | 0.50 | qPCR | 4.74 |
| OGL53 | 25 | Nahant, MA | July 12, 2016 | Asterias forbesi | Pyloric caeca | Healthy | NA | 4.5 | 2 mL | S23257 | 44.30 | Zymo Tissue & Insect | 0.75 | qPCR | 5.02 |
| OGL54 | 25 | Nahant, MA | July 12, 2016 | Asterias forbesi | Gonads | Healthy | NA | 4.5 | 2 mL | S23257 | 62.70 | Zymo Tissue & Insect | 49.80 | qPCR | 3.24 |
| OGL55 | 25 | Nahant, MA | July 12, 2016 | Asterias forbesi | Body Wall | Healthy | NA | 4.5 | 2 mL | S23257 | 68.90 | Zymo Tissue & Insect | 1.08 | qPCR | 3.19 |
| OGL56 | 26 | Nahant, MA | July 12, 2016 | Asterias forbesi | Whole animal | Healthy | NA | 3.1 | 2 mL | S23258 | 74.10 | Zymo Tissue & Insect | 0.56 | qPCR | 4.12 |
| OGL57 | 27 | Nahant, MA | July 12, 2016 | Asterias forbesi | Whole animal | Healthy | NA | 3.7 | 2 mL | S23259 | 70.80 | Zymo Tissue & Insect | 0.70 | qPCR | 4.18 |
| OGL58 | 28 | Nahant, MA | July 12, 2016 | Asterias forbesi | Whole animal | Healthy | NA | 3.1 | 2 mL | S23260 | 62.00 | Zymo Tissue & Insect | 0.49 | qPCR | 3.51 |
| OGL59 | 29 | Nahant, MA | July 12, 2016 | Asterias forbesi | Whole animal | Healthy | NA | 2.9 | 2 mL | S23261 | 62.00 | Zymo Tissue & Insect | 0.56 | qPCR | 4.34 |
| OGL60 | 30 | Nahant, MA | July 12, 2016 | Asterias forbesi | Whole animal | Healthy | NA | 3.4 | 2 mL | S23262 | 79.50 | Zymo Tissue & Insect | 0.56 | qPCR | 3.74 |
| OGL61 | 31 | Nahant, MA | July 12, 2016 | Asterias forbesi | Pyloric caeca | Healthy | NA | 5.2 | 2 mL | S23263 | 44.60 | Zymo Tissue & Insect | 1.10 | qPCR | 4.39 |
| OGL62 | 31 | Nahant, MA | July 12, 2016 | Asterias forbesi | Gonads | Healthy | NA | 5.2 | 2 mL | S23263 | 64.80 | Zymo Tissue & Insect | 52.19 | qPCR | 2.86 |
| OGL63 | 31 | Nahant, MA | July 12, 2016 | Asterias forbesi | Body Wall | Healthy | NA | 5.2 | 2 mL | S23263 | 75.60 | Zymo Tissue & Insect | 0.58 | qPCR | 2.71 |
| OGL64 | 32 | Nahant, MA | July 12, 2016 | Asterias forbesi | Whole animal | Healthy | NA | 3.6 | 2 mL | S23264 | 81.10 | Zymo Tissue & Insect | 4.15 | qPCR | 3.78 |
| OGL65 | 33 | Nahant, MA | July 12, 2016 | Asterias forbesi | Pyloric caeca | Healthy | NA | 5 | 2 mL | S23265 | 35.00 | Zymo Tissue & Insect | 1.41 | qPCR | 5.03 |
| OGL66 | 33 | Nahant, MA | July 12, 2016 | Asterias forbesi | Gonads | Healthy | NA | 5 | 2 mL | S23265 | 69.40 | Zymo Tissue & Insect | 54.81 | qPCR | 2.97 |
| OGL67 | 33 | Nahant, MA | July 12, 2016 | Asterias forbesi | Body Wall | Healthy | NA | 5 | 2 mL | S23265 | 70.20 | Zymo Tissue & Insect | 0.61 | qPCR | 3.29 |
| OGL68 | 34 | Nahant, MA | July 12, 2016 | Asterias forbesi | Pyloric caeca | Healthy | NA | 4.5 | 2 mL | S23266 | 54.50 | Zymo Tissue & Insect | 0.95 | qPCR | 4.63 |
| OGL70 | 34 | Nahant, MA | July 12, 2016 | Asterias forbesi | Body Wall | Healthy | NA | 4.5 | 2 mL | S23266 | 66.90 | Zymo Tissue & Insect | 0.53 | qPCR | 3.11 |
| OGL71 | 35 | Nahant, MA | July 12, 2016 | Asterias forbesi | Pyloric caeca | Healthy | NA | 8 | 2 mL | S23267 | 77.60 | Zymo Tissue & Insect | 0.54 | qPCR | 3.91 |
| OGL72 | 35 | Nahant, MA | July 12, 2016 | Asterias forbesi | Gonads | Healthy | NA | 8 | 2 mL | A27013 | 78.00 | Zymo Tissue & Insect | 53.03 | qPCR | 2.11 |
| OGL73 | 35 | Nahant, MA | July 15, 2016 | Asterias forbesi | Body Wall | Healthy | NA | 8 | 2 mL | S23267 | 71.40 | Zymo Tissue & Insect | 0.61 | qPCR | 2.12 |
| OGL74 | 36 | Nahant, MA | July 15, 2016 | Asterias forbesi | Pyloric caeca | Healthy | NA | 10.5 | 2 mL | S23268 | 82.90 | Zymo Tissue & Insect | 1.22 | qPCR | 3.83 |
| OGL75 | 36 | Nahant, MA | July 15, 2016 | Asterias forbesi | Gonads | Healthy | NA | 10.5 | 2 mL | S23268 | 67.40 | Zymo Tissue & Insect | 0.82 | qPCR | 2.38 |
| OGL76 | 36 | Nahant, MA | July 15, 2016 | Asterias forbesi | Body Wall | Healthy | NA | 10.5 | 2 mL | S23268 | 92.10 | Zymo Tissue & Insect | 0.64 | qPCR | 2.90 |
| OGL77 | 37 | Nahant, MA | July 15, 2016 | Asterias forbesi | Pyloric caeca | Healthy | NA | 7.8 | 2 mL | S23269 | 78.00 | Zymo Tissue & Insect | 2.54 | qPCR | 4.94 |
| OGL78 | 37 | Nahant, MA | July 15, 2016 | Asterias forbesi | Gonads | Healthy | NA | 7.8 | 2 mL | S23269 | 96.20 | Zymo Tissue & Insect | 1.20 | qPCR | 3.21 |
| OGL79 | 37 | Nahant, MA | July 15, 2016 | Asterias forbesi | Body Wall | Healthy | NA | 7.8 | 2 mL | S23269 | 73.00 | Zymo Tissue & Insect | 0.60 | qPCR | 3.01 |
| OGL80 | 38 | Nahant, MA | July 15, 2016 | Asterias forbesi | Pyloric caeca | Healthy | NA | 6.5 | 2 mL | S23270 | 33.40 | Zymo Tissue & Insect | 0.80 | qPCR | 3.74 |
| OGL81 | 38 | Nahant, MA | July 15, 2016 | Asterias forbesi | Gonads | Healthy | NA | 6.5 | 2 mL | S23270 | 68.00 | Zymo Tissue & Insect | 0.87 | qPCR | 1.69 |
| OGL82 | 38 | Nahant, MA | July 15, 2016 | Asterias forbesi | Body Wall | Healthy | NA | 6.5 | 2 mL | S23270 | 67.20 | Zymo Tissue & Insect | 0.61 | qPCR | 1.75 |
| OGL86 | 39 | Nahant, MA | July 15, 2016 | Asterias forbesi | Pyloric caeca | Healthy | NA | 7.8 | 2 mL | S23271 | 68.40 | Zymo Tissue & Insect | 0.69 | qPCR | 2.60 |
| OGL87 | 39 | Nahant, MA | July 15, 2016 | Asterias forbesi | Gonads | Healthy | NA | 7.8 | 2 mL | S23271 | 69.50 | Zymo Tissue & Insect | 0.50 | qPCR | 0.00 |
| OGL88 | 39 | Nahant, MA | July 15, 2016 | Asterias forbesi | Body Wall | Healthy | NA | 7.8 | 2 mL | S23271 | 85.20 | Zymo Tissue & Insect | 0.40 | qPCR | 0.00 |
| OGL89 | 40 | Nahant, MA | July 15, 2016 | Asterias forbesi | Pyloric caeca | Healthy | NA | 6.6 | 2 mL | S23272 | 58.00 | Zymo Tissue & Insect | 0.46 | qPCR | 3.89 |
| OGL90 | 40 | Nahant, MA | July 15, 2016 | Asterias forbesi | Gonads | Healthy | NA | 6.6 | 2 mL | S23272 | 80.00 | Zymo Tissue & Insect | 38.14 | qPCR | 1.76 |
| OGL91 | 40 | Nahant, MA | July 15, 2016 | Asterias forbesi | Body Wall | Healthy | NA | 6.6 | 2 mL | S23272 | 52.90 | Zymo Tissue & Insect | 0.35 | qPCR | 1.98 |
| OGL92 | 41 | Nahant, MA | July 15, 2016 | Asterias forbesi | Pyloric caeca | Healthy | NA | 4 | 2 mL | S23273 | 30.20 | Zymo Tissue & Insect | 0.48 | qPCR | 3.58 |
| OGL94 | 41 | Nahant, MA | July 15, 2016 | Asterias forbesi | Body Wall | Healthy | NA | 4 | 2 mL | S23273 | 53.20 | Zymo Tissue & Insect | 0.85 | qPCR | 2.46 |
| AF 1 BW | 1 | Bar Harbor, ME | May 20, 2017 | Asterias forbesi | Body Wall | Healthy | NA | NA | 2 mL | NA | 100.00 | Zymo Tissue & Insect | NA | qPCR | 0.00 |
| AF 1 CF | 1 | Bar Harbor, ME | May 20, 2017 | Asterias forbesi | Coelomic Fluid | Healthy | NA | NA | 2 mL | NA | 1000.00 | Zymo Tissue & Insect | 36.07 | qPCR | 0.00 |
| AF 1 G | 1 | Bar Harbor, ME | May 20, 2017 | Asterias forbesi | Gonads | Healthy | NA | NA | 2 mL | NA | 195.90 | Zymo Tissue & Insect | NA | qPCR | 0.00 |
| AF 1 PC | 1 | Bar Harbor, ME | May 20, 2017 | Asterias forbesi | Pyloric caeca | Healthy | NA | NA | 2 mL | NA | 132.20 | Zymo Tissue & Insect | NA | qPCR | 0.00 |
| AF 2 BW | 2 | Bar Harbor, ME | May 20, 2017 | Asterias forbesi | Body Wall | Healthy | NA | NA | 2 mL | NA | 86.30 | Zymo Tissue & Insect | 42.60 | qPCR | 0.00 |
| AF 2 CF | 2 | Bar Harbor, ME | May 20, 2017 | Asterias forbesi | Coelomic Fluid | Healthy | NA | NA | 2 mL | NA | 1000.00 | Zymo Tissue & Insect | 43.98 | qPCR | 0.00 |
| AF 2 G | 2 | Bar Harbor, ME | May 20, 2017 | Asterias forbesi | Gonads | Healthy | NA | NA | 2 mL | NA | 100.90 | Zymo Tissue & Insect | 41.11 | qPCR | 0.00 |
| AF 2 PC | 2 | Bar Harbor, ME | May 20, 2017 | Asterias forbesi | Pyloric caeca | Healthy | NA | NA | 2 mL | NA | 81.80 | Zymo Tissue & Insect | 37.83 | qPCR | 0.00 |
| AF 3 BW | 3 | Bar Harbor, ME | May 20, 2017 | Asterias forbesi | Body Wall | Healthy | NA | NA | 2 mL | NA | 162.60 | Zymo Tissue & Insect | 45.57 | qPCR | 0.00 |
| AF 3 CF | 3 | Bar Harbor, ME | May 20, 2017 | Asterias forbesi | Coelomic Fluid | Healthy | NA | NA | 2 mL | NA | 1000.00 | Zymo Tissue & Insect | 32.76 | qPCR | 0.00 |
| AF 3 G | 3 | Bar Harbor, ME | May 20, 2017 | Asterias forbesi | Gonads | Healthy | NA | NA | 2 mL | NA | 75.10 | Zymo Tissue & Insect | 39.55 | qPCR | 0.00 |
| AF 3 PC | 3 | Bar Harbor, ME | May 20, 2017 | Asterias forbesi | Pyloric caeca | Healthy | NA | NA | 2 mL | NA | 79.90 | Zymo Tissue & Insect | 40.28 | qPCR | 0.00 |

| | | | | | | | | | | | | | | | |
|---|---|---|---|---|---|---|---|---|---|---|---|---|---|---|---|
| AF 4 BW | 4 | Bar Harbor, ME | May 20, 2017 | Asterias forbesi | Body Wall | Healthy | NA | NA | 2 mL | NA | 126.60 | Zymo Tissue & Insect | 5.67 | qPCR | 0.00 |
| AF 4 CF | 4 | Bar Harbor, ME | May 20, 2017 | Asterias forbesi | Coelomic Fluid | Healthy | NA | NA | 2 mL | NA | 1000.00 | Zymo Tissue & Insect | 0.46 | qPCR | 0.00 |
| AF 4 G | 4 | Bar Harbor, ME | May 20, 2017 | Asterias forbesi | Gonads | Healthy | NA | NA | 2 mL | NA | 43.10 | Zymo Tissue & Insect | 0.81 | qPCR | 0.00 |
| AF 4 PC | 4 | Bar Harbor, ME | May 20, 2017 | Asterias forbesi | Pyloric caeca | Healthy | NA | NA | 2 mL | NA | 63.40 | Zymo Tissue & Insect | 9.88 | qPCR | 0.00 |
| AF 5 BW | 5 | Bar Harbor, ME | May 20, 2017 | Asterias forbesi | Body Wall | Healthy | NA | NA | 2 mL | NA | 84.60 | Zymo Tissue & Insect | 7.09 | qPCR | 0.00 |
| AF 5 CF | 5 | Bar Harbor, ME | May 20, 2017 | Asterias forbesi | Coelomic Fluid | Healthy | NA | NA | 2 mL | NA | 1000.00 | Zymo Tissue & Insect | 1.16 | qPCR | 0.00 |
| AF 5 G | 5 | Bar Harbor, ME | May 20, 2017 | Asterias forbesi | Gonads | Healthy | NA | NA | 2 mL | NA | 70.50 | Zymo Tissue & Insect | 30.18 | qPCR | 0.00 |
| AF 5 PC | 5 | Bar Harbor, ME | May 20, 2017 | Asterias forbesi | Pyloric caeca | Healthy | NA | NA | 2 mL | NA | 72.20 | Zymo Tissue & Insect | 36.21 | qPCR | 0.00 |
| AF 6 BW | 6 | Bar Harbor, ME | May 20, 2017 | Asterias forbesi | Body Wall | Healthy | NA | NA | 2 mL | NA | 64.30 | Zymo Tissue & Insect | 3.92 | qPCR | 0.00 |
| AF 6 CF | 6 | Bar Harbor, ME | May 20, 2017 | Asterias forbesi | Coelomic Fluid | Healthy | NA | NA | 2 mL | NA | 1000.00 | Zymo Tissue & Insect | 0.83 | qPCR | 0.00 |
| AF 6 G | 6 | Bar Harbor, ME | May 20, 2017 | Asterias forbesi | Gonads | Healthy | NA | NA | 2 mL | NA | 45.60 | Zymo Tissue & Insect | 2.85 | qPCR | 0.00 |
| AF 6 PC | 6 | Bar Harbor, ME | May 20, 2017 | Asterias forbesi | Pyloric caeca | Healthy | NA | 18 | 2 mL | NA | 72.60 | Zymo Tissue & Insect | 28.00 | qPCR | 0.00 |
| MBL01 | 1 | Woods Hole, MA | May 5, 2019 | Asterias forbesi | Pyloric caeca | Healthy | NA | 18 | 2 mL | NA | 45.00 | Zymo Quick DNA Miniprep Plus | 0.52 | PCR | NA |
| MBL02 | 2 | Woods Hole, MA | May 5, 2019 | Asterias forbesi | Pyloric caeca | Healthy | NA | 17 | 2 mL | NA | 61.80 | Zymo Quick DNA Miniprep Plus | 0.41 | PCR | NA |
| MBL03 | 3 | Woods Hole, MA | May 5, 2019 | Asterias forbesi | Pyloric caeca | Healthy | NA | 18.5 | 2 mL | NA | 19.40 | Zymo Quick DNA Miniprep Plus | 33.64 | PCR | NA |
| MBL04 | 4 | Woods Hole, MA | May 5, 2019 | Asterias forbesi | Pyloric caeca | Healthy | NA | 18.5 | 2 mL | NA | 24.60 | Zymo Quick DNA Miniprep Plus | 32.81 | PCR | NA |
| MBL05 | 5 | Woods Hole, MA | May 5, 2019 | Asterias forbesi | Pyloric caeca | Healthy | NA | 18.5 | 2 mL | NA | 47.70 | Zymo Quick DNA Miniprep Plus | 0.53 | PCR | NA |
| MBL26 G | 6 | Woods Hole, MA | June 22, 2019 | Asterias forbesi | Oocytes | Healthy | Female | 19 | 2 mL | NA | NA | Zymo Quick DNA Miniprep Plus | 23.99 | PCR | NA |
| MBL26 PC | 6 | Woods Hole, MA | June 22, 2019 | Asterias forbesi | Pyloric caeca | Healthy | Female | 19 | 2 mL | NA | NA | Zymo Quick DNA Miniprep Plus | 1.72 | PCR | NA |
| MBL27 G | 7 | Woods Hole, MA | June 22, 2019 | Asterias forbesi | Oocytes | Healthy | Female | 18 | 2 mL | NA | NA | Zymo Quick DNA Miniprep Plus | 5.63 | PCR | NA |
| MBL27 PC | 7 | Woods Hole, MA | June 22, 2019 | Asterias forbesi | Pyloric caeca | Healthy | Female | 18 | 2 mL | NA | NA | Zymo Quick DNA Miniprep Plus | 0.33 | PCR | NA |
| MBL28 G | 8 | Woods Hole, MA | June 22, 2019 | Asterias forbesi | Gonads | Healthy | Male | 15 | 2 mL | NA | NA | Zymo Quick DNA Miniprep Plus | 13.77 | PCR | NA |
| MBL29 G | 8 | Woods Hole, MA | June 22, 2019 | Asterias forbesi | Gonads | Healthy | Male | 14 | 2 mL | NA | NA | Zymo Quick DNA Miniprep Plus | 38.84 | PCR | NA |
| MBL30 G | 9 | Woods Hole, MA | June 22, 2019 | Asterias forbesi | Gonads | Healthy | Male | 14 | 2 mL | NA | NA | Zymo Quick DNA Miniprep Plus | 38.19 | PCR | NA |
| MBL31 G | 9 | Woods Hole, MA | June 22, 2019 | Asterias forbesi | Oocytes | Healthy | Female | 14.4 | 2 mL | NA | NA | Zymo Quick DNA Miniprep Plus | 3.61 | PCR | NA |
| MBL31 PC | 10 | Woods Hole, MA | June 22, 2019 | Asterias forbesi | Pyloric caeca | Healthy | Female | 14.4 | 2 mL | NA | NA | Zymo Quick DNA Miniprep Plus | 0.18 | PCR | NA |
| MBL32 G | 10 | Woods Hole, MA | June 22, 2019 | Asterias forbesi | Gonads | Healthy | Male | 13 | 2 mL | NA | NA | Zymo Quick DNA Miniprep Plus | 38.47 | PCR | NA |
| MBL33 G | 11 | Woods Hole, MA | June 22, 2019 | Asterias forbesi | Oocytes | Healthy | Female | 20 | 2 mL | NA | NA | Zymo Quick DNA Miniprep Plus | 14.65 | PCR | NA |
| MBL33 PC | 11 | Woods Hole, MA | June 22, 2019 | Asterias forbesi | Pyloric caeca | Healthy | Female | 20 | 2 mL | NA | NA | Zymo Quick DNA Miniprep Plus | 2.43 | PCR | NA |
| MBL34 G | 12 | Woods Hole, MA | June 22, 2019 | Asterias forbesi | Oocytes | Healthy | Female | 15 | 2 mL | NA | NA | Zymo Quick DNA Miniprep Plus | 4.47 | PCR | NA |
| MBL34 PC | 12 | Woods Hole, MA | June 22, 2019 | Asterias forbesi | Pyloric caeca | Healthy | Female | 15 | 2 mL | NA | NA | Zymo Quick DNA Miniprep Plus | 1.02 | PCR | NA |
| MBL35 G | 13 | Woods Hole, MA | June 22, 2019 | Asterias forbesi | Oocytes | Healthy | Female | 18.2 | 2 mL | NA | NA | Zymo Quick DNA Miniprep Plus | 6.38 | PCR | NA |
| MBL35 PC | 13 | Woods Hole, MA | June 22, 2019 | Asterias forbesi | Pyloric caeca | Healthy | Female | 18.2 | 2 mL | NA | NA | Zymo Quick DNA Miniprep Plus | 0.42 | PCR | NA |
| MBL36 G | 14 | Woods Hole, MA | June 29, 2019 | Asterias forbesi | Oocytes | Healthy | Female | 15.6 | 2 mL | NA | NA | Zymo Quick DNA Miniprep Plus | 20.53 | PCR | NA |
| MBL36 PC | 14 | Woods Hole, MA | June 29, 2019 | Asterias forbesi | Pyloric caeca | Healthy | Female | 15.6 | 2 mL | NA | NA | Zymo Quick DNA Miniprep Plus | 0.45 | PCR | NA |
| MBL37 G | 15 | Woods Hole, MA | June 29, 2019 | Asterias forbesi | Oocytes | Healthy | Female | 16.8 | 2 mL | NA | NA | Zymo Quick DNA Miniprep Plus | 21.26 | PCR | NA |
| MBL37 PC | 15 | Woods Hole, MA | June 29, 2019 | Asterias forbesi | Pyloric caeca | Healthy | Female | 16.8 | 2 mL | NA | NA | Zymo Quick DNA Miniprep Plus | 0.35 | PCR | NA |
| MBL38 G | 16 | Woods Hole, MA | June 29, 2019 | Asterias forbesi | Oocytes | Healthy | Female | 16.8 | 2 mL | NA | NA | Zymo Quick DNA Miniprep Plus | 26.93 | PCR | NA |
| MBL38 PC | 16 | Woods Hole, MA | June 29, 2019 | Asterias forbesi | Pyloric caeca | Healthy | Female | 16.8 | 2 mL | NA | NA | Zymo Quick DNA Miniprep Plus | 0.59 | PCR | NA |
| MBL39 G | 17 | Woods Hole, MA | June 29, 2019 | Asterias forbesi | Gonads | Healthy | Male | 18 | 2 mL | NA | NA | Zymo Quick DNA Miniprep Plus | 1.47 | PCR | NA |
| MBL40 G | 18 | Woods Hole, MA | June 29, 2019 | Asterias forbesi | Gonads | Healthy | Male | 19 | 2 mL | NA | NA | Zymo Quick DNA Miniprep Plus | 6.69 | PCR | NA |
| MBL41 G | 19 | Woods Hole, MA | June 29, 2019 | Asterias forbesi | Gonads | Healthy | Male | 18.6 | 2 mL | NA | NA | Zymo Quick DNA Miniprep Plus | 18.33 | PCR | NA |
| MBL42 G | 20 | Woods Hole, MA | June 29, 2019 | Asterias forbesi | Gonads | Healthy | Male | 23 | 2 mL | NA | NA | Zymo Quick DNA Miniprep Plus | 33.40 | PCR | NA |
| MBL43 G | 21 | Woods Hole, MA | June 29, 2019 | Asterias forbesi | Oocytes | Healthy | Female | 18 | 2 mL | NA | NA | Zymo Quick DNA Miniprep Plus | 17.49 | PCR | NA |
| MBL44 G | 22 | Woods Hole, MA | June 29, 2019 | Asterias forbesi | Oocytes | Healthy | Female | 21.2 | 2 mL | NA | NA | Zymo Quick DNA Miniprep Plus | 13.07 | PCR | NA |
| MBL45 G | 23 | Woods Hole, MA | June 29, 2019 | Asterias forbesi | Oocytes | Healthy | Female | 20.8 | 2 mL | NA | NA | Zymo Quick DNA Miniprep Plus | 13.32 | PCR | NA |
| MBL46 G | 24 | Woods Hole, MA | June 29, 2019 | Asterias forbesi | Oocytes | Healthy | Female | 24 | 2 mL | NA | NA | Zymo Quick DNA Miniprep Plus | 25.91 | PCR | NA |
| MBL47 G | 25 | Woods Hole, MA | June 29, 2019 | Asterias forbesi | Gonads | Healthy | Male | 17.6 | 2 mL | NA | NA | Zymo Quick DNA Miniprep Plus | 35.43 | PCR | NA |
| MBL48 G | 26 | Woods Hole, MA | July 8, 2019 | Asterias forbesi | Oocytes | Healthy | Female | 23 | 2 mL | NA | NA | Zymo Quick DNA Miniprep Plus | 2.60 | PCR | NA |
| MBL49 G | 27 | Woods Hole, MA | July 8, 2019 | Asterias forbesi | Gonads | Healthy | Male | 17 | 2 mL | NA | NA | Zymo Quick DNA Miniprep Plus | 33.78 | PCR | NA |

| | | | | | | | | | | | | | | | |
|---|---|---|---|---|---|---|---|---|---|---|---|---|---|---|---|
| MBL50 G | 28 | Woods Hole, MA | July 8, 2019 | Asterias forbesi | Oocytes | Healthy | Female | 12 | 2 mL | NA | NA | Zymo Quick DNA Miniprep Plus | 5.06 | PCR | NA |
| MBL50 PC | 28 | Woods Hole, MA | July 8, 2019 | Asterias forbesi | Pyloric caeca | Healthy | Female | 12 | 2 mL | NA | NA | Zymo Quick DNA Miniprep Plus | 0.40 | PCR | NA |
| MBL51 G | 29 | Woods Hole, MA | July 8, 2019 | Asterias forbesi | Gonads | Healthy | Male | 18 | 2 mL | NA | NA | Zymo Quick DNA Miniprep Plus | 35.39 | PCR | NA |
| MBL52 G | 29 | Woods Hole, MA | July 8, 2019 | Asterias forbesi | Gonads | Healthy | Male | 10 | 2 mL | NA | NA | Zymo Quick DNA Miniprep Plus | 36.04 | PCR | NA |
| MYA01 | 1 | Mystic Aquarium, CT | October 12, 2012 | Asterias forbesi | Pyloric caeca | Diseased | NA | NA | 2 mL | NA | 81.60 | Zymo Tissue & Insect | 0.38 | qPCR | 2.43 |
| MYA02 | 1 | Mystic Aquarium, CT | October 12, 2012 | Asterias forbesi | Gonads | Diseased | NA | NA | 2 mL | NA | 62.10 | Zymo Tissue & Insect | 0.43 | qPCR | 3.03 |
| MYA03 | 1 | Mystic Aquarium, CT | October 12, 2012 | Asterias forbesi | Body Wall | Diseased | NA | NA | 2 mL | NA | 71.10 | Zymo Tissue & Insect | 0.28 | qPCR | 2.76 |
| MYA04 | 2 | Mystic Aquarium, CT | October 12, 2012 | Asterias forbesi | Pyloric caeca | Diseased | NA | NA | 2 mL | NA | 65.60 | Zymo Tissue & Insect | 0.63 | qPCR | 3.08 |
| MYA05 | 2 | Mystic Aquarium, CT | October 12, 2012 | Asterias forbesi | Body Wall | Diseased | NA | NA | 2 mL | NA | 87.70 | Zymo Tissue & Insect | 0.31 | qPCR | 2.72 |
| MYA06 | 3 | Mystic Aquarium, CT | October 16, 2012 | Asterias forbesi | Pyloric caeca | Diseased | NA | NA | 2 mL | NA | 101.90 | Zymo Tissue & Insect | 0.58 | qPCR | 3.14 |
| MYA07 | 3 | Mystic Aquarium, CT | October 16, 2012 | Asterias forbesi | Gonads | Diseased | NA | NA | 2 mL | NA | 77.30 | Zymo Tissue & Insect | 1.04 | qPCR | 2.52 |
| MYA08 | 3 | Mystic Aquarium, CT | October 16, 2012 | Asterias forbesi | Body Wall | Diseased | NA | NA | 2 mL | NA | 61.80 | Zymo Tissue & Insect | 0.34 | qPCR | 2.92 |
| MYA09 | 4 | Mystic Aquarium, CT | October 25, 2012 | Asterias forbesi | Pyloric caeca | Diseased | NA | NA | 2 mL | NA | 77.30 | Zymo Tissue & Insect | 0.42 | qPCR | 2.69 |
| MYA10 | 4 | Mystic Aquarium, CT | October 25, 2012 | Asterias forbesi | Gonads | Diseased | NA | NA | 2 mL | NA | 74.90 | Zymo Tissue & Insect | 0.36 | qPCR | 2.76 |
| MYA11 | 4 | Mystic Aquarium, CT | October 25, 2012 | Asterias forbesi | Body Wall | Diseased | NA | NA | 2 mL | NA | 80.60 | Zymo Tissue & Insect | 25.66 | qPCR | 2.35 |
| MYA12 | 5 | Mystic Aquarium, CT | October 22, 2012 | Asterias forbesi | Pyloric caeca | Diseased | NA | NA | 2 mL | NA | 65.30 | Zymo Tissue & Insect | 0.82 | qPCR | 2.84 |
| MYA13 | 5 | Mystic Aquarium, CT | October 22, 2012 | Asterias forbesi | Gonads | Diseased | NA | NA | 2 mL | NA | 22.50 | Zymo Tissue & Insect | 0.36 | qPCR | 1.89 |
| MYA14 | 5 | Mystic Aquarium, CT | October 22, 2012 | Asterias forbesi | Body Wall | Diseased | NA | NA | 2 mL | NA | 74.80 | Zymo Tissue & Insect | 0.25 | qPCR | 1.69 |
| MYA15 | 6 | Mystic Aquarium, CT | October 13, 2012 | Asterias forbesi | Pyloric caeca | Diseased | NA | NA | 2 mL | NA | 65.60 | Zymo Tissue & Insect | 0.57 | qPCR | 1.98 |
| MYA16 | 6 | Mystic Aquarium, CT | October 13, 2012 | Asterias forbesi | Gonads | Diseased | NA | NA | 2 mL | NA | 14.80 | Zymo Tissue & Insect | 0.27 | qPCR | 3.06 |
| MYA17 | 6 | Mystic Aquarium, CT | October 13, 2012 | Asterias forbesi | Body Wall | Diseased | NA | NA | 2 mL | NA | 85.40 | Zymo Tissue & Insect | 0.32 | qPCR | 1.65 |
| MYA18 | 7 | Mystic Aquarium, CT | October 15, 2012 | Asterias forbesi | Pyloric caeca | Diseased | NA | NA | 2 mL | NA | 63.30 | Zymo Tissue & Insect | 1.16 | qPCR | 1.92 |
| MYA19 | 7 | Mystic Aquarium, CT | October 15, 2012 | Asterias forbesi | Gonads | Diseased | NA | NA | 2 mL | NA | 66.00 | Zymo Tissue & Insect | 31.54 | qPCR | 2.00 |
| MYA20 | 7 | Mystic Aquarium, CT | October 15, 2012 | Asterias forbesi | Body Wall | Diseased | NA | NA | 2 mL | NA | 81.80 | Zymo Tissue & Insect | 0.39 | qPCR | 1.87 |
| MYA21 | 8 | Mystic Aquarium, CT | October 15, 2012 | Asterias forbesi | Pyloric caeca | Diseased | NA | NA | 2 mL | NA | 66.10 | Zymo Tissue & Insect | 0.36 | qPCR | 2.31 |
| MYA22 | 8 | Mystic Aquarium, CT | October 15, 2012 | Asterias forbesi | Gonads | Diseased | NA | NA | 2 mL | NA | 15.90 | Zymo Tissue & Insect | 0.63 | qPCR | 2.39 |
| MYA23 | 8 | Mystic Aquarium, CT | October 15, 2012 | Asterias forbesi | Body Wall | Diseased | NA | NA | 2 mL | NA | 64.90 | Zymo Tissue & Insect | 0.31 | qPCR, PCR | 3.74 |
| MYA24 | 9 | Mystic Aquarium, CT | October 15, 2012 | Asterias forbesi | Pyloric caeca | Diseased | NA | NA | 2 mL | NA | 66.50 | Zymo Tissue & Insect | 0.77 | qPCR | 2.12 |
| MYA25 | 9 | Mystic Aquarium, CT | October 15, 2012 | Asterias forbesi | Gonads | Diseased | NA | NA | 2 mL | NA | 62.60 | Zymo Tissue & Insect | 0.54 | qPCR | 0.00 |
| MYA26 | 9 | Mystic Aquarium, CT | October 15, 2012 | Asterias forbesi | Body Wall | Diseased | NA | NA | 2 mL | NA | 91.40 | Zymo Tissue & Insect | 0.40 | qPCR | 1.79 |
| MYA27 | 10 | Mystic Aquarium, CT | October 21, 2012 | Asterias forbesi | Pyloric caeca | Diseased | NA | NA | 2 mL | NA | 63.00 | Zymo Tissue & Insect | 0.27 | qPCR | 2.51 |
| MYA29 | 10 | Mystic Aquarium, CT | October 21, 2012 | Asterias forbesi | Body Wall | Diseased | NA | NA | 2 mL | NA | 81.00 | Zymo Tissue & Insect | 0.28 | qPCR | 2.40 |
| MYA30 | 11 | Mystic Aquarium, CT | October 21, 2012 | Asterias forbesi | Pyloric caeca | Diseased | NA | NA | 2 mL | NA | 62.50 | Zymo Tissue & Insect | 0.35 | qPCR, PCR | 3.61 |
| MYA32 | 11 | Mystic Aquarium, CT | October 21, 2012 | Asterias forbesi | Body Wall | Diseased | NA | NA | 2 mL | NA | 81.10 | Zymo Tissue & Insect | 0.21 | qPCR | 2.55 |
| MYA33 | 12 | Mystic Aquarium, CT | October 21, 2012 | Asterias forbesi | Pyloric caeca | Diseased | NA | NA | 2 mL | NA | 64.30 | Zymo Tissue & Insect | 0.32 | qPCR | 2.58 |
| MYA34 | 12 | Mystic Aquarium, CT | October 21, 2012 | Asterias forbesi | Gonads | Diseased | NA | NA | 2 mL | NA | 35.60 | Zymo Tissue & Insect | 0.74 | qPCR | 2.88 |
| MYA35 | 12 | Mystic Aquarium, CT | October 21, 2012 | Asterias forbesi | Body Wall | Diseased | NA | NA | 2 mL | NA | 85.30 | Zymo Tissue & Insect | 0.43 | qPCR | 1.98 |
| MYA36 | 13 | Mystic Aquarium, CT | October 16, 2012 | Asterias forbesi | Pyloric caeca | Diseased | NA | NA | 2 mL | NA | 88.10 | Zymo Tissue & Insect | 0.55 | qPCR | 1.91 |
| MYA37 | 13 | Mystic Aquarium, CT | October 16, 2012 | Asterias forbesi | Gonads | Diseased | NA | NA | 2 mL | NA | 69.20 | Zymo Tissue & Insect | 3.44 | qPCR | 2.03 |
| MYA38 | 13 | Mystic Aquarium, CT | October 16, 2012 | Asterias forbesi | Body Wall | Diseased | NA | NA | 2 mL | NA | 85.10 | Zymo Tissue & Insect | 0.24 | qPCR | 2.22 |
| MYA39 | 14 | Mystic Aquarium, CT | October 16, 2012 | Asterias forbesi | Pyloric caeca | Diseased | NA | NA | 2 mL | NA | 62.50 | Zymo Tissue & Insect | 0.55 | qPCR | 2.72 |
| MYA41 | 14 | Mystic Aquarium, CT | October 16, 2012 | Asterias forbesi | Body Wall | Diseased | NA | NA | 2 mL | NA | 76.80 | Zymo Tissue & Insect | 0.29 | qPCR | 2.91 |
| MYA42 | 15 | Mystic Aquarium, CT | October 16, 2012 | Asterias forbesi | Pyloric caeca | Diseased | NA | NA | 2 mL | NA | 68.00 | Zymo Tissue & Insect | 39.36 | qPCR | 0.00 |
| MYA44 | 15 | Mystic Aquarium, CT | October 16, 2012 | Asterias forbesi | Body Wall | Diseased | NA | NA | 2 mL | NA | 67.20 | Zymo Tissue & Insect | 21.20 | qPCR | 0.00 |
| MYA45 | 16 | Mystic Aquarium, CT | October 16, 2012 | Asterias forbesi | Pyloric caeca | Diseased | NA | NA | 2 mL | NA | 83.70 | Zymo Tissue & Insect | 0.43 | qPCR | 0.00 |
| MYA46 | 16 | Mystic Aquarium, CT | October 16, 2012 | Asterias forbesi | Gonads | Diseased | NA | NA | 2 mL | NA | 81.60 | Zymo Tissue & Insect | 0.54 | qPCR | 0.00 |
| MYA47 | 16 | Mystic Aquarium, CT | October 16, 2012 | Asterias forbesi | Body Wall | Diseased | NA | NA | 2 mL | NA | 66.40 | Zymo Tissue & Insect | 0.35 | qPCR | 0.00 |
| MYA48 | 17 | Mystic Aquarium, CT | October 20, 2012 | Asterias forbesi | Pyloric caeca | Diseased | NA | NA | 2 mL | NA | 69.30 | Zymo Tissue & Insect | 0.34 | qPCR | 0.00 |
| MYA49 | 17 | Mystic Aquarium, CT | October 20, 2012 | Asterias forbesi | Gonads | Diseased | NA | NA | 2 mL | NA | 75.50 | Zymo Tissue & Insect | 2.52 | qPCR | 0.00 |
| MYA50 | 17 | Mystic Aquarium, CT | October 20, 2012 | Asterias forbesi | Body Wall | Diseased | NA | NA | 2 mL | NA | 89.70 | Zymo Tissue & Insect | 1.74 | qPCR | 0.00 |

| | | | | | | | | | | | | | | | |
|---|---|---|---|---|---|---|---|---|---|---|---|---|---|---|---|
| MYA51 | 18 | Mystic Aquarium, CT | July 25, 2014 | Asterias forbesi | Pyloric caeca | Diseased | NA | NA | 2 mL | NA | 83.30 | Zymo Tissue & Insect | 0.45 | qPCR | 0.00 |
| MYA52 | 18 | Mystic Aquarium, CT | July 25, 2014 | Asterias forbesi | Gonads | Diseased | NA | NA | 2 mL | NA | 65.80 | Zymo Tissue & Insect | 11.18 | qPCR | 0.00 |
| MYA53 | 18 | Mystic Aquarium, CT | July 25, 2014 | Asterias forbesi | Body Wall | Diseased | NA | NA | 2 mL | NA | 58.00 | Zymo Tissue & Insect | 29.66 | qPCR | 0.00 |
| MYA54 | 19 | Mystic Aquarium, CT | October 13, 2012 | Asterias forbesi | Pyloric caeca | Diseased | NA | NA | 2 mL | NA | 64.70 | Zymo Tissue & Insect | 0.39 | qPCR | 0.00 |
| MYA55 | 19 | Mystic Aquarium, CT | October 13, 2012 | Asterias forbesi | Gonads | Diseased | NA | NA | 2 mL | NA | 74.20 | Zymo Tissue & Insect | 0.33 | qPCR | 0.00 |
| MYA56 | 19 | Mystic Aquarium, CT | October 13, 2012 | Asterias forbesi | Body Wall | Diseased | NA | NA | 2 mL | NA | 71.00 | Zymo Tissue & Insect | 0.99 | qPCR | 1.88 |
| MYA57 | 20 | Mystic Aquarium, CT | October 25, 2013 | Asterias forbesi | Pyloric caeca | Diseased | NA | NA | 2 mL | NA | 81.10 | Zymo Tissue & Insect | 3.05 | qPCR | 0.00 |
| MYA58 | 20 | Mystic Aquarium, CT | October 25, 2013 | Asterias forbesi | Gonads | Diseased | NA | NA | 2 mL | NA | 46.00 | Zymo Tissue & Insect | 0.40 | qPCR | 0.00 |
| MYA59 | 20 | Mystic Aquarium, CT | October 25, 2013 | Asterias forbesi | Body Wall | Diseased | NA | NA | 2 mL | NA | 74.40 | Zymo Tissue & Insect | 1.81 | qPCR | 0.00 |
| MYA60 | 21 | Mystic Aquarium, CT | February 23, 2014 | Asterias forbesi | Pyloric caeca | Diseased | NA | NA | 2 mL | NA | 70.60 | Zymo Tissue & Insect | 46.14 | qPCR | 0.00 |
| MYA61 | 21 | Mystic Aquarium, CT | February 23, 2014 | Asterias forbesi | Gonads | Diseased | NA | NA | 2 mL | NA | 74.00 | Zymo Tissue & Insect | 0.47 | qPCR | 0.00 |
| MYA62 | 21 | Mystic Aquarium, CT | February 23, 2014 | Asterias forbesi | Body Wall | Diseased | NA | NA | 2 mL | NA | 60.50 | Zymo Tissue & Insect | 57.34 | qPCR | 0.00 |
| SML1 | 1 | Shoals Marine Lab, ME | June 25, 2016 | Asterias forbesi | Coelomic Fluid | Healthy | NA | 8 | 2 mL | NA | 900.00 | Zymo Tissue & Insect | 0.72 | qPCR | 0.00 |
| SML100 | 22 | Shoals Marine Lab, ME | July 9, 2016 | Henricia sp | Pyloric caeca | Healthy | NA | 4.3 | 2 mL | NA | 65.00 | Zymo Tissue & Insect | 0.26 | qPCR | 2.44 |
| SML101 | 22 | Shoals Marine Lab, ME | July 9, 2016 | Henricia sp | Body Wall | Healthy | NA | 4.3 | 2 mL | NA | 76.20 | Zymo Tissue & Insect | 0.42 | qPCR | 3.21 |
| SML102 | 23 | Shoals Marine Lab, ME | June 25, 2016 | Asterias rubens | Whole animal | Healthy | NA | 2 | 2 mL | NA | 70.20 | Zymo Tissue & Insect | 6.28 | qPCR | 3.42 |
| SML103 | 24 | Shoals Marine Lab, ME | July 9, 2016 | Asterias rubens | Coelomic Fluid | Healthy | NA | 6.5 | 2 mL | NA | 1000.00 | Zymo Tissue & Insect | 0.34 | qPCR | 0.00 |
| SML104 | 24 | Shoals Marine Lab, ME | July 9, 2016 | Asterias rubens | Pyloric caeca | Healthy | NA | 6.5 | 2 mL | NA | 70.10 | Zymo Tissue & Insect | 4.60 | qPCR, PCR | 3.16 |
| SML105 | 24 | Shoals Marine Lab, ME | July 9, 2016 | Asterias rubens | Body Wall | Healthy | NA | 6.5 | 2 mL | NA | 68.40 | Zymo Tissue & Insect | 0.81 | qPCR | 0.00 |
| SML107 | 25 | Shoals Marine Lab, ME | July 9, 2016 | Henricia sp | Pyloric caeca | Healthy | NA | 3.6 | 2 mL | NA | 65.40 | Zymo Tissue & Insect | 1.33 | qPCR, PCR | 5.21 |
| SML108 | 25 | Shoals Marine Lab, ME | July 9, 2016 | Henricia sp | Body Wall | Healthy | NA | 3.6 | 2 mL | NA | 63.00 | Zymo Tissue & Insect | 1.40 | qPCR | 3.36 |
| SML109 | 26 | Shoals Marine Lab, ME | July 9, 2016 | Henricia sp | Pyloric caeca | Healthy | NA | 4 | 2 mL | NA | 79.30 | Zymo Tissue & Insect | 0.56 | qPCR, PCR | 4.50 |
| SML110 | 26 | Shoals Marine Lab, ME | July 9, 2016 | Henricia sp | Body Wall | Healthy | NA | 4 | 2 mL | NA | 64.80 | Zymo Tissue & Insect | 0.49 | qPCR | 3.07 |
| SML111 | 27 | Shoals Marine Lab, ME | July 9, 2016 | Henricia sp | Pyloric caeca | Healthy | NA | 3 | 2 mL | NA | 65.60 | Zymo Tissue & Insect | 2.52 | qPCR | 3.71 |
| SML112 | 27 | Shoals Marine Lab, ME | July 9, 2016 | Henricia sp | Body Wall | Healthy | NA | 3 | 2 mL | NA | 72.20 | Zymo Tissue & Insect | 0.28 | qPCR, PCR | 4.26 |
| SML113 | 28 | Shoals Marine Lab, ME | July 9, 2016 | Henricia sp | Pyloric caeca | Healthy | NA | 4.5 | 2 mL | NA | 74.70 | Zymo Tissue & Insect | 0.35 | qPCR | 3.73 |
| SML114 | 28 | Shoals Marine Lab, ME | July 9, 2016 | Henricia sp | Gonads | Healthy | NA | 4.5 | 2 mL | NA | 72.70 | Zymo Tissue & Insect | 24.79 | qPCR | 0.00 |
| SML115 | 28 | Shoals Marine Lab, ME | July 9, 2016 | Henricia sp | Body Wall | Healthy | NA | 4.5 | 2 mL | NA | 74.60 | Zymo Tissue & Insect | 50.05 | qPCR | 2.23 |
| SML116 | 29 | Shoals Marine Lab, ME | July 9, 2016 | Henricia sp | Whole animal | Healthy | NA | 3 | 2 mL | NA | 65.40 | Zymo Tissue & Insect | 0.56 | qPCR | 3.33 |
| SML117 | 30 | Shoals Marine Lab, ME | July 9, 2016 | Henricia sp | Pyloric caeca | Healthy | NA | 3.4 | 2 mL | NA | 21.80 | Zymo Tissue & Insect | 1.02 | qPCR, PCR | 4.14 |
| SML118 | 30 | Shoals Marine Lab, ME | July 9, 2016 | Henricia sp | Gonads | Healthy | NA | 3.4 | 2 mL | NA | 22.20 | Zymo Tissue & Insect | 41.73 | qPCR | 2.51 |
| SML119 | 31 | Shoals Marine Lab, ME | July 9, 2016 | Henricia sp | Coelomic Fluid | Healthy | NA | 7.5 | 2 mL | NA | 150.00 | Zymo Tissue & Insect | 0.40 | qPCR | 0.00 |
| SML120 | 31 | Shoals Marine Lab, ME | July 9, 2016 | Henricia sp | Pyloric caeca | Healthy | NA | 7.5 | 2 mL | NA | 69.30 | Zymo Tissue & Insect | 23.41 | qPCR | 2.90 |
| SML121 | 31 | Shoals Marine Lab, ME | July 9, 2016 | Henricia sp | Gonads | Healthy | NA | 7.5 | 2 mL | NA | 75.20 | Zymo Tissue & Insect | 47.95 | qPCR | 0.00 |
| SML122 | 31 | Shoals Marine Lab, ME | July 9, 2016 | Henricia sp | Body Wall | Healthy | NA | 7.5 | 2 mL | NA | 80.00 | Zymo Tissue & Insect | 0.35 | qPCR | 1.87 |
| SML123 | 30 | Shoals Marine Lab, ME | July 9, 2016 | Henricia sp | Body Wall | Healthy | NA | 3.4 | 2 mL | NA | 61.60 | Zymo Tissue & Insect | 50.27 | qPCR | 2.94 |
| SML124 | 32 | Shoals Marine Lab, ME | July 9, 2016 | Asterias rubens | Coelomic Fluid | Healthy | NA | 10.5 | 2 mL | NA | 800.00 | Zymo Tissue & Insect | 0.43 | qPCR | 0.00 |
| SML125 | 32 | Shoals Marine Lab, ME | July 9, 2016 | Asterias rubens | Pyloric caeca | Healthy | NA | 10.5 | 2 mL | NA | 80.00 | Zymo Tissue & Insect | 1.84 | qPCR, PCR | 3.43 |
| SML126 | 32 | Shoals Marine Lab, ME | July 9, 2016 | Asterias rubens | Gonads | Healthy | NA | 10.5 | 2 mL | NA | 78.40 | Zymo Tissue & Insect | 37.79 | qPCR | 1.89 |
| SML127 | 32 | Shoals Marine Lab, ME | July 9, 2016 | Asterias rubens | Body Wall | Healthy | NA | 10.5 | 2 mL | NA | 74.50 | Zymo Tissue & Insect | 0.45 | qPCR | 0.00 |
| SML128 | 33 | Shoals Marine Lab, ME | July 9, 2016 | Asterias rubens | Coelomic Fluid | Healthy | NA | 8 | 2 mL | NA | 750.00 | Zymo Tissue & Insect | 0.51 | qPCR | 0.00 |
| SML129 | 33 | Shoals Marine Lab, ME | July 9, 2016 | Asterias rubens | Pyloric caeca | Healthy | NA | 8 | 2 mL | NA | 750.00 | Zymo Tissue & Insect | 4.47 | qPCR, PCR | 3.11 |
| SML13 | 3 | Shoals Marine Lab, ME | June 25, 2016 | Asterias rubens | Coelomic Fluid | Healthy | NA | 8.5 | 2 mL | NA | 400.00 | Zymo Tissue & Insect | 0.86 | qPCR | 0.00 |
| SML130 | 33 | Shoals Marine Lab, ME | July 9, 2016 | Asterias rubens | Gonads | Healthy | NA | 8 | 2 mL | NA | 48.00 | Zymo Tissue & Insect | 1.31 | qPCR | 0.00 |
| SML131 | 33 | Shoals Marine Lab, ME | July 9, 2016 | Asterias rubens | Body Wall | Healthy | NA | 8 | 2 mL | NA | 74.50 | Zymo Tissue & Insect | 0.47 | qPCR | 0.00 |
| SML132 | 34 | Shoals Marine Lab, ME | July 9, 2016 | Asterias rubens | Coelomic Fluid | Healthy | NA | 7 | 2 mL | NA | 1000.00 | Zymo Tissue & Insect | 0.57 | qPCR | 0.00 |
| SML133 | 34 | Shoals Marine Lab, ME | July 9, 2016 | Asterias rubens | Pyloric caeca | Healthy | NA | 7 | 2 mL | NA | 69.00 | Zymo Tissue & Insect | 7.16 | qPCR | 3.84 |
| SML134 | 34 | Shoals Marine Lab, ME | July 9, 2016 | Asterias rubens | Gonads | Healthy | NA | 7 | 2 mL | NA | 24.00 | Zymo Tissue & Insect | 8.84 | qPCR | 0.00 |
| SML135 | 34 | Shoals Marine Lab, ME | July 9, 2016 | Asterias rubens | Body Wall | Healthy | NA | 7 | 2 mL | NA | 69.90 | Zymo Tissue & Insect | 0.40 | qPCR | 0.00 |
| SML137 | 35 | Shoals Marine Lab, ME | July 9, 2016 | Asterias rubens | Pyloric caeca | Healthy | NA | 8.5 | 2 mL | NA | 71.80 | Zymo Tissue & Insect | 5.40 | qPCR | 0.00 |

| | | | | | | | | | | | | | | | |
|---|---|---|---|---|---|---|---|---|---|---|---|---|---|---|---|
| SML138 | 35 | Shoals Marine Lab, ME | July 9, 2016 | Asterias rubens | Gonads | Healthy | NA | 8.5 | 2 mL | NA | 73.70 | Zymo Tissue & Insect | 3.32 | qPCR | 0.00 |
| SML139 | 35 | Shoals Marine Lab, ME | July 9, 2016 | Asterias rubens | Body Wall | Healthy | NA | 8.5 | 2 mL | NA | 64.30 | Zymo Tissue & Insect | 2.19 | qPCR | 1.76 |
| SML14 | 3 | Shoals Marine Lab, ME | June 25, 2016 | Asterias rubens | Pyloric caeca | Healthy | NA | 8.5 | 2 mL | NA | 79.00 | Zymo Tissue & Insect | 0.65 | qPCR | 2.45 |
| SML140 | 36 | Shoals Marine Lab, ME | July 9, 2016 | Asterias rubens | Pyloric caeca | Healthy | NA | 7 | 2 mL | NA | 66.80 | Zymo Tissue & Insect | 0.34 | qPCR | 3.21 |
| SML141 | 36 | Shoals Marine Lab, ME | July 9, 2016 | Asterias rubens | Body Wall | Healthy | NA | 7 | 2 mL | NA | 73.10 | Zymo Tissue & Insect | 0.39 | qPCR | 0.00 |
| SML142 | 37 | Shoals Marine Lab, ME | July 9, 2016 | Asterias rubens | Coelomic Fluid | Healthy | NA | 8.5 | 2 mL | NA | 1000.00 | Zymo Tissue & Insect | 1.28 | qPCR | 0.00 |
| SML143 | 37 | Shoals Marine Lab, ME | July 9, 2016 | Asterias rubens | Pyloric caeca | Healthy | NA | 8.5 | 2 mL | NA | 78.30 | Zymo Tissue & Insect | 3.60 | qPCR | 2.74 |
| SML144 | 37 | Shoals Marine Lab, ME | July 9, 2016 | Asterias rubens | Gonads | Healthy | NA | 8.5 | 2 mL | NA | 15.20 | Zymo Tissue & Insect | 40.43 | qPCR | 0.00 |
| SML145 | 37 | Shoals Marine Lab, ME | July 9, 2016 | Asterias rubens | Body Wall | Healthy | NA | 8.5 | 2 mL | NA | 68.00 | Zymo Tissue & Insect | 0.39 | qPCR | 0.00 |
| SML146 | 38 | Shoals Marine Lab, ME | July 9, 2016 | Asterias rubens | Coelomic Fluid | Healthy | NA | 6.5 | 2 mL | NA | 1000.00 | Zymo Tissue & Insect | 0.34 | qPCR | 0.00 |
| SML147 | 38 | Shoals Marine Lab, ME | July 9, 2016 | Asterias rubens | Pyloric caeca | Healthy | NA | 6.5 | 2 mL | NA | 74.90 | Zymo Tissue & Insect | 8.57 | qPCR | 2.54 |
| SML148 | 38 | Shoals Marine Lab, ME | July 9, 2016 | Asterias rubens | Body Wall | Healthy | NA | 6.5 | 2 mL | NA | 63.60 | Zymo Tissue & Insect | 0.44 | qPCR | 0.00 |
| SML15 | 3 | Shoals Marine Lab, ME | June 25, 2016 | Asterias rubens | Gonads | Healthy | NA | 8.5 | 2 mL | NA | 65.50 | Zymo Tissue & Insect | 0.42 | qPCR | 2.11 |
| SML150 | 39 | Shoals Marine Lab, ME | July 9, 2016 | Asterias rubens | Coelomic Fluid | Healthy | NA | 8 | 2 mL | NA | 400.00 | Zymo Tissue & Insect | 1.26 | qPCR | 0.00 |
| SML151 | 39 | Shoals Marine Lab, ME | July 9, 2016 | Asterias rubens | Pyloric caeca | Healthy | NA | 8 | 2 mL | NA | 76.60 | Zymo Tissue & Insect | 2.11 | qPCR | 3.03 |
| SML152 | 39 | Shoals Marine Lab, ME | July 9, 2016 | Asterias rubens | Gonads | Healthy | NA | 8 | 2 mL | NA | 66.20 | Zymo Tissue & Insect | 3.06 | qPCR | 0.00 |
| SML153 | 39 | Shoals Marine Lab, ME | July 9, 2016 | Asterias rubens | Body Wall | Healthy | NA | 8 | 2 mL | NA | 64.50 | Zymo Tissue & Insect | 0.92 | qPCR | 1.82 |
| SML154 | 40 | Shoals Marine Lab, ME | July 9, 2016 | Asterias rubens | Pyloric caeca | Healthy | NA | 5.5 | 2 mL | NA | 68.70 | Zymo Tissue & Insect | 7.54 | qPCR | 3.11 |
| SML155 | 40 | Shoals Marine Lab, ME | July 9, 2016 | Asterias rubens | Body Wall | Healthy | NA | 5.5 | 2 mL | NA | 73.90 | Zymo Tissue & Insect | 1.06 | qPCR | 1.72 |
| SML157 | 41 | Shoals Marine Lab, ME | July 9, 2016 | Asterias rubens | Coelomic Fluid | Healthy | NA | 6.2 | 2 mL | NA | 700.00 | Zymo Tissue & Insect | 1.22 | qPCR | 0.00 |
| SML158 | 41 | Shoals Marine Lab, ME | July 9, 2016 | Asterias rubens | Pyloric caeca | Healthy | NA | 6.2 | 2 mL | NA | 63.10 | Zymo Tissue & Insect | 4.59 | qPCR | 3.13 |
| SML159 | 41 | Shoals Marine Lab, ME | July 9, 2016 | Asterias rubens | Gonads | Healthy | NA | 6.2 | 2 mL | NA | 41.90 | Zymo Tissue & Insect | 36.85 | qPCR | 0.00 |
| SML16 | 3 | Shoals Marine Lab, ME | June 25, 2016 | Asterias rubens | Body Wall | Healthy | NA | 8.5 | 2 mL | NA | 75.50 | Zymo Tissue & Insect | 0.50 | qPCR | 1.83 |
| SML160 | 41 | Shoals Marine Lab, ME | July 9, 2016 | Asterias rubens | Body Wall | Healthy | NA | 6.2 | 2 mL | NA | 69.60 | Zymo Tissue & Insect | 1.96 | qPCR | 1.98 |
| SML161 | 42 | Shoals Marine Lab, ME | July 9, 2016 | Asterias rubens | Coelomic Fluid | Healthy | NA | 7 | 2 mL | NA | 400.00 | Zymo Tissue & Insect | 8.13 | qPCR | 0.00 |
| SML162 | 42 | Shoals Marine Lab, ME | July 9, 2016 | Asterias rubens | Pyloric caeca | Healthy | NA | 7 | 2 mL | NA | 63.30 | Zymo Tissue & Insect | 13.75 | qPCR | 3.24 |
| SML163 | 42 | Shoals Marine Lab, ME | July 9, 2016 | Asterias rubens | Gonads | Healthy | NA | 7 | 2 mL | NA | 74.40 | Zymo Tissue & Insect | 46.73 | qPCR | 0.00 |
| SML164 | 42 | Shoals Marine Lab, ME | July 9, 2016 | Asterias rubens | Body Wall | Healthy | NA | 7 | 2 mL | NA | 73.50 | Zymo Tissue & Insect | 0.56 | qPCR | 3.09 |
| SML165 | 43 | Shoals Marine Lab, ME | July 9, 2016 | Asterias rubens | Coelomic Fluid | Healthy | NA | 7 | 2 mL | NA | 500.00 | Zymo Tissue & Insect | 1.16 | qPCR | 0.00 |
| SML166 | 43 | Shoals Marine Lab, ME | July 9, 2016 | Asterias rubens | Pyloric caeca | Healthy | NA | 7 | 2 mL | NA | 70.10 | Zymo Tissue & Insect | 3.89 | qPCR, PCR | 4.00 |
| SML167 | 43 | Shoals Marine Lab, ME | July 9, 2016 | Asterias rubens | Gonads | Healthy | NA | 7 | 2 mL | NA | 48.30 | Zymo Tissue & Insect | 4.58 | qPCR | 2.23 |
| SML168 | 43 | Shoals Marine Lab, ME | July 9, 2016 | Asterias rubens | Body Wall | Healthy | NA | 7 | 2 mL | NA | 79.60 | Zymo Tissue & Insect | 0.57 | qPCR | 2.23 |
| SML169 | 44 | Shoals Marine Lab, ME | July 9, 2016 | Asterias rubens | Coelomic Fluid | Healthy | NA | 6.2 | 2 mL | NA | 700.00 | Zymo Tissue & Insect | 1.48 | qPCR | 0.00 |
| SML170 | 44 | Shoals Marine Lab, ME | July 9, 2016 | Asterias rubens | Pyloric caeca | Healthy | NA | 6.2 | 2 mL | NA | 85.60 | Zymo Tissue & Insect | 8.92 | qPCR, PCR | 2.95 |
| SML171 | 44 | Shoals Marine Lab, ME | July 9, 2016 | Asterias rubens | Body Wall | Healthy | NA | 6.2 | 2 mL | NA | 66.40 | Zymo Tissue & Insect | 3.34 | qPCR | 1.98 |
| SML173 | 45 | Shoals Marine Lab, ME | July 9, 2016 | Asterias rubens | Coelomic Fluid | Healthy | NA | 7 | 2 mL | NA | 500.00 | Zymo Tissue & Insect | 0.41 | qPCR | 0.00 |
| SML174 | 45 | Shoals Marine Lab, ME | July 9, 2016 | Asterias rubens | Pyloric caeca | Healthy | NA | 7 | 2 mL | NA | 77.30 | Zymo Tissue & Insect | 8.39 | qPCR | 2.96 |
| SML175 | 45 | Shoals Marine Lab, ME | July 9, 2016 | Asterias rubens | Gonads | Healthy | NA | 7 | 2 mL | NA | 65.40 | Zymo Tissue & Insect | 23.25 | qPCR | 0.00 |
| SML176 | 45 | Shoals Marine Lab, ME | July 9, 2016 | Asterias rubens | Body Wall | Healthy | NA | 7 | 2 mL | NA | 76.80 | Zymo Tissue & Insect | 15.65 | qPCR | 3.11 |
| SML177 | 46 | Shoals Marine Lab, ME | June 25, 2016 | Asterias rubens | Coelomic Fluid | Healthy | NA | 8.2 | 2 mL | NA | 800.00 | Zymo Tissue & Insect | 2.45 | qPCR | 0.00 |
| SML178 | 46 | Shoals Marine Lab, ME | June 25, 2016 | Asterias rubens | Pyloric caeca | Healthy | NA | 8.2 | 2 mL | NA | 77.40 | Zymo Tissue & Insect | 1.93 | qPCR | 2.94 |
| SML180 | 46 | Shoals Marine Lab, ME | June 25, 2016 | Asterias rubens | Body Wall | Healthy | NA | 8.2 | 2 mL | NA | 59.30 | Zymo Tissue & Insect | 1.93 | qPCR | 2.37 |
| SML19 | 4 | Shoals Marine Lab, ME | June 25, 2016 | Henricia sp | Coelomic Fluid | Healthy | NA | 10.5 | 2 mL | NA | 1000.00 | Zymo Tissue & Insect | 3.51 | qPCR | 0.00 |
| SML196 | 47 | Shoals Marine Lab, ME | June 25, 2016 | Asterias rubens | Coelomic Fluid | Healthy | NA | 8.5 | 2 mL | NA | 400.00 | Zymo Tissue & Insect | 0.69 | qPCR | 0.00 |
| SML197 | 47 | Shoals Marine Lab, ME | June 25, 2016 | Asterias rubens | Pyloric caeca | Healthy | NA | 8.5 | 2 mL | NA | 66.80 | Zymo Tissue & Insect | 4.65 | qPCR | 2.76 |
| SML198 | 47 | Shoals Marine Lab, ME | June 25, 2016 | Asterias rubens | Gonads | Healthy | NA | 8.5 | 2 mL | NA | 53.70 | Zymo Tissue & Insect | 2.35 | qPCR | 1.76 |
| SML199 | 47 | Shoals Marine Lab, ME | June 25, 2016 | Asterias rubens | Body Wall | Healthy | NA | 8.5 | 2 mL | NA | 61.70 | Zymo Tissue & Insect | 1.08 | qPCR | 2.15 |
| SML2 | 1 | Shoals Marine Lab, ME | June 25, 2016 | Asterias forbesi | Pyloric caeca | Healthy | NA | 8 | 2 mL | NA | 64.90 | Zymo Tissue & Insect | 22.16 | qPCR, PCR | 3.19 |
| SML20 | 4 | Shoals Marine Lab, ME | June 25, 2016 | Henricia sp | Pyloric caeca | Healthy | NA | 10.5 | 2 mL | NA | 71.20 | Zymo Tissue & Insect | 29.76 | qPCR, PCR | 2.55 |
| SML200 | 48 | Shoals Marine Lab, ME | June 25, 2016 | Asterias rubens | Coelomic Fluid | Healthy | NA | 8.5 | 2 mL | NA | 800.00 | Zymo Tissue & Insect | 1.71 | qPCR | 0.00 |

| | | | | | | | | | | | | | | | |
|---|---|---|---|---|---|---|---|---|---|---|---|---|---|---|---|
| SML201 | 48 | Shoals Marine Lab, ME | June 25, 2016 | Asterias rubens | Pyloric caeca | Healthy | NA | 8.5 | 2 mL | NA | 64.00 | Zymo Tissue & Insect | 16.20 | qPCR | 2.56 |
| SML202 | 48 | Shoals Marine Lab, ME | June 25, 2016 | Asterias rubens | Body Wall | Healthy | NA | 8.5 | 2 mL | NA | 85.70 | Zymo Tissue & Insect | 13.01 | qPCR | 2.06 |
| SML205 | 49 | Shoals Marine Lab, ME | June 25, 2016 | Asterias rubens | Coelomic Fluid | Diseased | NA | 10.7 | 2 mL | NA | 1000.00 | Zymo Tissue & Insect | 0.47 | qPCR | 0.00 |
| SML206 | 49 | Shoals Marine Lab, ME | June 25, 2016 | Asterias rubens | Pyloric caeca | Diseased | NA | 10.7 | 2 mL | NA | 77.00 | Zymo Tissue & Insect | 4.66 | qPCR | 2.82 |
| SML207 | 49 | Shoals Marine Lab, ME | June 25, 2016 | Asterias rubens | Gonads | Diseased | NA | 10.7 | 2 mL | NA | 68.90 | Zymo Tissue & Insect | 1.07 | qPCR | 1.86 |
| SML208 | 49 | Shoals Marine Lab, ME | June 25, 2016 | Asterias rubens | Body Wall | Diseased | NA | 10.7 | 2 mL | NA | 63.50 | Zymo Tissue & Insect | 4.14 | qPCR | 2.31 |
| SML209 | 50 | Shoals Marine Lab, ME | June 25, 2016 | Asterias rubens | Coelomic Fluid | Healthy | NA | 7.5 | 2 mL | NA | 500.00 | Zymo Tissue & Insect | 0.38 | qPCR | 0.00 |
| SML210 | 50 | Shoals Marine Lab, ME | June 25, 2016 | Asterias rubens | Pyloric caeca | Healthy | NA | 7.5 | 2 mL | NA | 73.90 | Zymo Tissue & Insect | 13.40 | qPCR | 2.50 |
| SML211 | 50 | Shoals Marine Lab, ME | June 25, 2016 | Asterias rubens | Body Wall | Healthy | NA | 7.5 | 2 mL | NA | 62.70 | Zymo Tissue & Insect | 2.60 | qPCR | 2.24 |
| SML212 | 51 | Shoals Marine Lab, ME | June 25, 2016 | Asterias rubens | Coelomic Fluid | Healthy | NA | 7 | 2 mL | NA | 1000.00 | Zymo Tissue & Insect | 7.26 | qPCR | 0.00 |
| SML213 | 51 | Shoals Marine Lab, ME | June 25, 2016 | Asterias rubens | Pyloric caeca | Healthy | NA | 7 | 2 mL | NA | 73.70 | Zymo Tissue & Insect | 5.60 | qPCR | 2.82 |
| SML214 | 51 | Shoals Marine Lab, ME | June 25, 2016 | Asterias rubens | Body Wall | Healthy | NA | 7 | 2 mL | NA | 82.50 | Zymo Tissue & Insect | 0.58 | qPCR | 1.64 |
| SML22 | 4 | Shoals Marine Lab, ME | June 25, 2016 | Henricia sp | Gonads | Healthy | NA | 10.5 | 2 mL | NA | 64.60 | Zymo Tissue & Insect | 38.70 | qPCR | 0.00 |
| SML24 | 4 | Shoals Marine Lab, ME | June 25, 2016 | Henricia sp | Body Wall | Healthy | NA | 10.5 | 2 mL | NA | 78.90 | Zymo Tissue & Insect | 1.28 | qPCR | 2.10 |
| SML29 | 5 | Shoals Marine Lab, ME | June 25, 2016 | Henricia sp | Body Wall | Diseased | NA | 3.4 | 2 mL | NA | 66.00 | Zymo Tissue & Insect | 0.42 | qPCR | 3.08 |
| SML33 | 6 | Shoals Marine Lab, ME | June 25, 2016 | Asterias rubens | Coelomic Fluid | Healthy | NA | 5.2 | 2 mL | NA | 140.00 | Zymo Tissue & Insect | 0.44 | qPCR | 0.00 |
| SML34 | 6 | Shoals Marine Lab, ME | June 25, 2016 | Asterias rubens | Pyloric caeca | Healthy | NA | 5.2 | 2 mL | NA | 60.00 | Zymo Tissue & Insect | 32.62 | qPCR, PCR | 2.45 |
| SML35 | 6 | Shoals Marine Lab, ME | June 25, 2016 | Asterias rubens | Gonads | Healthy | NA | 5.2 | 2 mL | NA | 37.20 | Zymo Tissue & Insect | 4.60 | qPCR | 1.95 |
| SML36 | 6 | Shoals Marine Lab, ME | June 25, 2016 | Asterias rubens | Body Wall | Healthy | NA | 5.2 | 2 mL | NA | 80.00 | Zymo Tissue & Insect | 0.65 | qPCR | 1.81 |
| SML39 | 7 | Shoals Marine Lab, ME | June 25, 2016 | Asterias rubens | Coelomic Fluid | Healthy | NA | 6.2 | 2 mL | NA | 150.00 | Zymo Tissue & Insect | 0.36 | qPCR | 0.00 |
| SML40 | 7 | Shoals Marine Lab, ME | June 25, 2016 | Asterias rubens | Pyloric caeca | Healthy | NA | 6.2 | 2 mL | NA | 66.50 | Zymo Tissue & Insect | 7.31 | qPCR | 2.51 |
| SML42 | 7 | Shoals Marine Lab, ME | June 25, 2016 | Asterias rubens | Body Wall | Healthy | NA | 6.2 | 2 mL | NA | 66.50 | Zymo Tissue & Insect | 0.73 | qPCR | 1.60 |
| SML45 | 8 | Shoals Marine Lab, ME | June 25, 2016 | Asterias forbesi | Coelomic Fluid | Healthy | NA | 9.5 | 2 mL | NA | 1000.00 | Zymo Tissue & Insect | 14.11 | qPCR | 0.00 |
| SML46 | 8 | Shoals Marine Lab, ME | June 25, 2016 | Asterias forbesi | Pyloric caeca | Healthy | NA | 9.5 | 2 mL | NA | 67.50 | Zymo Tissue & Insect | 0.29 | qPCR, PCR | 3.49 |
| SML47 | 8 | Shoals Marine Lab, ME | June 25, 2016 | Asterias forbesi | Gonads | Healthy | NA | 9.5 | 2 mL | NA | 79.20 | Zymo Tissue & Insect | 47.40 | qPCR | 0.00 |
| SML48 | 8 | Shoals Marine Lab, ME | June 25, 2016 | Asterias forbesi | Body Wall | Healthy | NA | 9.5 | 2 mL | NA | 79.00 | Zymo Tissue & Insect | 0.70 | qPCR, PCR | 1.94 |
| SML5 | 2 | Shoals Marine Lab, ME | June 25, 2016 | Asterias rubens | Coelomic Fluid | Healthy | NA | 7 | 2 mL | NA | 400.00 | Zymo Tissue & Insect | 1.22 | qPCR | 0.00 |
| SML51 | 9 | Shoals Marine Lab, ME | June 25, 2016 | Asterias rubens | Coelomic Fluid | Healthy | NA | 8.3 | 2 mL | NA | 180.00 | Zymo Tissue & Insect | 0.57 | qPCR | 0.00 |
| SML52 | 9 | Shoals Marine Lab, ME | June 25, 2016 | Asterias rubens | Pyloric caeca | Healthy | NA | 8.3 | 2 mL | NA | 64.80 | Zymo Tissue & Insect | 7.46 | qPCR | 2.04 |
| SML53 | 9 | Shoals Marine Lab, ME | June 25, 2016 | Asterias rubens | Gonads | Healthy | NA | 8.3 | 2 mL | NA | 72.70 | Zymo Tissue & Insect | 9.54 | qPCR | 1.73 |
| SML54 | 9 | Shoals Marine Lab, ME | June 25, 2016 | Asterias rubens | Body Wall | Healthy | NA | 8.3 | 2 mL | NA | 73.20 | Zymo Tissue & Insect | 2.67 | qPCR | 1.96 |
| SML57 | 10 | Shoals Marine Lab, ME | June 25, 2016 | Asterias rubens | Body Wall | Healthy | NA | 5.4 | 2 mL | NA | 87.10 | Zymo Tissue & Insect | 15.18 | qPCR | 2.32 |
| SML6 | 2 | Shoals Marine Lab, ME | June 25, 2016 | Asterias rubens | Pyloric caeca | Healthy | NA | 7 | 2 mL | NA | 78.10 | Zymo Tissue & Insect | 9.09 | qPCR, PCR | 3.34 |
| SML63 | 11 | Shoals Marine Lab, ME | June 28, 2016 | Henricia sp | Pyloric caeca | Diseased | NA | 14 | 2 mL | NA | 69.20 | Zymo Tissue & Insect | 0.43 | qPCR, PCR | 2.36 |
| SML64 | 11 | Shoals Marine Lab, ME | June 28, 2016 | Henricia sp | Gonads | Diseased | NA | 14 | 2 mL | NA | 78.90 | Zymo Tissue & Insect | 0.32 | qPCR | 2.17 |
| SML65 | 11 | Shoals Marine Lab, ME | June 28, 2016 | Henricia sp | Body Wall | Diseased | NA | 14 | 2 mL | NA | 75.80 | Zymo Tissue & Insect | 0.37 | qPCR | 2.19 |
| SML67 | 12 | Shoals Marine Lab, ME | June 28, 2016 | Henricia sp | Pyloric caeca | Diseased | NA | 12 | 2 mL | NA | 73.10 | Zymo Tissue & Insect | 10.61 | qPCR | 2.78 |
| SML68 | 12 | Shoals Marine Lab, ME | June 28, 2016 | Henricia sp | Gonads | Diseased | NA | 12 | 2 mL | NA | 73.20 | Zymo Tissue & Insect | 40.66 | qPCR | 0.00 |
| SML69 | 12 | Shoals Marine Lab, ME | June 28, 2016 | Henricia sp | Body Wall | Diseased | NA | 12 | 2 mL | NA | 69.90 | Zymo Tissue & Insect | 0.35 | qPCR | 2.09 |
| SML7 | 2 | Shoals Marine Lab, ME | June 25, 2016 | Asterias rubens | Gonads | Healthy | NA | 7 | 2 mL | NA | 33.10 | Zymo Tissue & Insect | 1.58 | qPCR | 2.40 |
| SML72 | 13 | Shoals Marine Lab, ME | June 28, 2016 | Henricia sp | Pyloric caeca | Diseased | NA | 13 | 2 mL | NA | 78.30 | Zymo Tissue & Insect | 0.46 | qPCR, PCR | 2.97 |
| SML73 | 13 | Shoals Marine Lab, ME | June 28, 2016 | Henricia sp | Gonads | Diseased | NA | 13 | 2 mL | NA | 63.20 | Zymo Tissue & Insect | 0.49 | qPCR | 2.06 |
| SML74 | 13 | Shoals Marine Lab, ME | June 28, 2016 | Henricia sp | Body Wall | Diseased | NA | 13 | 2 mL | NA | 79.60 | Zymo Tissue & Insect | 0.36 | qPCR | 3.00 |
| SML75 | 14 | Shoals Marine Lab, ME | June 25, 2016 | Asterias rubens | Whole animal | Healthy | NA | 2.5 | 2 mL | NA | 76.10 | Zymo Tissue & Insect | 0.64 | qPCR | 2.05 |
| SML78 | 15 | Shoals Marine Lab, ME | June 25, 2016 | Asterias rubens | Coelomic Fluid | Healthy | NA | 6.6 | 2 mL | NA | 400.00 | Zymo Tissue & Insect | 0.39 | qPCR | 0.00 |
| SML79 | 15 | Shoals Marine Lab, ME | June 25, 2016 | Asterias rubens | Pyloric caeca | Healthy | NA | 6.6 | 2 mL | NA | 78.50 | Zymo Tissue & Insect | 9.94 | qPCR | 2.08 |
| SML8 | 2 | Shoals Marine Lab, ME | June 25, 2016 | Asterias rubens | Body Wall | Healthy | NA | 7 | 2 mL | NA | 72.10 | Zymo Tissue & Insect | 0.39 | qPCR | 1.64 |
| SML81 | 15 | Shoals Marine Lab, ME | June 25, 2016 | Asterias rubens | Body Wall | Healthy | NA | 6.6 | 2 mL | NA | 75.00 | Zymo Tissue & Insect | 0.85 | qPCR | 1.94 |
| SML82 | 16 | Shoals Marine Lab, ME | July 9, 2016 | Asterias rubens | Pyloric caeca | Healthy | NA | 5.5 | 2 mL | NA | 62.00 | Zymo Tissue & Insect | 0.70 | qPCR | 0.00 |
| SML86 | 17 | Shoals Marine Lab, ME | June 25, 2016 | Asterias rubens | Coelomic Fluid | Healthy | NA | 11.4 | 2 mL | NA | 1000.00 | Zymo Tissue & Insect | 0.44 | qPCR | 0.00 |

| | | | | | | | | | | | | | | | |
|---|---|---|---|---|---|---|---|---|---|---|---|---|---|---|---|
| SML87 | 17 | Shoals Marine Lab, ME | June 25, 2016 | Asterias rubens | Pyloric caeca | Healthy | NA | 11.4 | 2 mL | NA | 72.80 | Zymo Tissue & Insect | 4.52 | qPCR, PCR | 3.43 |
| SML88 | 17 | Shoals Marine Lab, ME | June 25, 2016 | Asterias rubens | Gonads | Healthy | NA | 11.4 | 2 mL | NA | 65.10 | Zymo Tissue & Insect | 1.87 | qPCR | 0.00 |
| SML89 | 17 | Shoals Marine Lab, ME | June 25, 2016 | Asterias rubens | Body Wall | Healthy | NA | 11.4 | 2 mL | NA | 69.40 | Zymo Tissue & Insect | 0.26 | qPCR | 2.46 |
| SML90 | 18 | Shoals Marine Lab, ME | July 9, 2016 | Asterias rubens | Pyloric caeca | Healthy | NA | 4 | 2 mL | NA | 69.20 | Zymo Tissue & Insect | 4.52 | qPCR, PCR | 3.01 |
| SML91 | 18 | Shoals Marine Lab, ME | July 9, 2016 | Asterias rubens | Body Wall | Healthy | NA | 4 | 2 mL | NA | 72.60 | Zymo Tissue & Insect | 0.41 | qPCR | 0.00 |
| SML92 | 19 | Shoals Marine Lab, ME | July 9, 2016 | Asterias rubens | Coelomic Fluid | Healthy | NA | 9 | 2 mL | NA | 500.00 | Zymo Tissue & Insect | 0.98 | qPCR | 0.00 |
| SML93 | 19 | Shoals Marine Lab, ME | July 9, 2016 | Asterias rubens | Pyloric caeca | Healthy | NA | 9 | 2 mL | NA | 66.20 | Zymo Tissue & Insect | 2.05 | qPCR | 1.94 |
| SML94 | 19 | Shoals Marine Lab, ME | July 9, 2016 | Asterias rubens | Gonads | Healthy | NA | 9 | 2 mL | NA | 41.00 | Zymo Tissue & Insect | 0.77 | qPCR | 0.00 |
| SML95 | 19 | Shoals Marine Lab, ME | July 9, 2016 | Asterias rubens | Body Wall | Healthy | NA | 9 | 2 mL | NA | 73.40 | Zymo Tissue & Insect | 0.26 | qPCR | 0.00 |
| SML96 | 20 | Shoals Marine Lab, ME | July 9, 2016 | Asterias rubens | Pyloric caeca | Healthy | NA | 5 | 2 mL | NA | 67.40 | Zymo Tissue & Insect | 2.32 | qPCR | 2.51 |
| SML97 | 20 | Shoals Marine Lab, ME | July 9, 2016 | Asterias rubens | Body Wall | Healthy | NA | 5 | 2 mL | NA | 76.90 | Zymo Tissue & Insect | 0.44 | qPCR | 0.00 |
| SML98 | 21 | Shoals Marine Lab, ME | July 9, 2016 | Asterias rubens | Pyloric caeca | Healthy | NA | 4 | 2 mL | NA | 64.00 | Zymo Tissue & Insect | 5.94 | qPCR, PCR | 3.35 |
| SML99 | 21 | Shoals Marine Lab, ME | July 9, 2016 | Asterias rubens | Body Wall | Healthy | NA | 4 | 2 mL | NA | 74.20 | Zymo Tissue & Insect | 0.29 | qPCR | 2.01 |
| AF1 | 42 | Nahant, MA | September 23, 2015 | Asterias forbesi | Whole animal | Diseased | NA | NA | 2 mL | S22798 | 65.00 | Zymo Tissue & Insect | NA | Metagenomics | NA |
| AF2 | 43 | Nahant, MA | September 23, 2015 | Asterias forbesi | Whole animal | Diseased | NA | NA | 2 mL | S22799 | 65.00 | Zymo Tissue & Insect | NA | Metagenomics | NA |
| AF3 | 44 | Nahant, MA | September 23, 2015 | Asterias forbesi | Whole animal | Diseased | NA | NA | 2 mL | S22800 | 65.00 | Zymo Tissue & Insect | NA | Metagenomics | NA |
| AF4 | 45 | Nahant, MA | September 23, 2015 | Asterias forbesi | Whole animal | Diseased | NA | NA | 2 mL | S22801 | 65.00 | Zymo Tissue & Insect | NA | Metagenomics | NA |
| AF5 | 46 | Nahant, MA | October 14, 2015 | Asterias forbesi | Whole animal | Diseased | NA | NA | 2 mL | S22802 | 65.00 | Zymo Tissue & Insect | NA | Metagenomics | NA |
| AF6 | 47 | Nahant, MA | October 14, 2015 | Asterias forbesi | Whole animal | Diseased | NA | NA | 2 mL | S22803 | 65.00 | Zymo Tissue & Insect | NA | Metagenomics | NA |
