## Supplemental Table 2 for "A highly prevalent and pervasive densovirus discovered among sea stars from the North American Atlantic Coast"

**Supplementary Table**
**qPCR primers/probe and PCR primers**

| Purpose | Sequence | |
|---|---|---|
| AfaDV<br><br>quantitative<br>(qPCR) | Standard | TTCGTAATAGCACCTTCGTCACCAGCTAAATATAGATTTTCTCC<br>AACTCTGATGAAACCAGCACGTCCACCAGACTGATGAATGCGC<br>AGTAATTGTCTA |
| | Internal Probe | [FAM] TGATGAAACCAGCACGTCCACCAGA [TAMRA] |
| | L-primer<br>position 5664 | CGTAATAGCACCTTCGTCACC |
| | R-primer<br>position 5738 | GACAATTACTGCGCATTCATCA |
| Thermocycling parameters (qPCR): 1 cycle of 50°C for 2 min followed by 95 °C for 2 min, 50 cycles of 95 °C for 15s and 58 °C for 1 min | | |
| AfaDV<br><br>genome<br>verification<br><br>(PCR) | L-primer – 1<br>position 243 | TTTGAGGTCATATGGGCGGA |
| | R-primer – 1<br>position 833 | CTTGTCACAACTCCTTTTCGC |
| | L-primer – 2<br>position 554 | ACGCCACTCAGTATGCAGTA |
| | R-primer – 2<br>position 1061 | TCCCAAGCTTTGCCAGAGTA |
| | L-primer – 3<br>position 831 | TGCGAAAAGGAGTTGTGACA |
| | R-primer – 3<br>position 1361 | TGCAAACGCTATCTTCTTCTCC |
| | L-primer – 4<br>position 1266 | TGCCGGATCTGACCATTGAT |
| | R-primer – 4<br>position 1702 | TTCTCGACATACCTGGAGCA |
| | L-primer - 5<br>Position 1520 | AAGCAGCAAAGACATGGAGC |
| | R-primer - 5<br>position 2075 | GATCCGGTTCGTCATCATCG |
| | L-primer - 6<br>position 1979 | GGAGAGCGGACTTGATGGAT |
| | R-primer – 6<br>position 2564 | AGAAATTCTTACCCGCTGAAGG |
| | L-primer – 7<br>position 2378 | GTGCAGGGTACGGTAATTTTG |
| | R-primer – 7<br>Position 2919 | ACAGCAAGCGGATTAGGTTTC |
| | L-primer – 8<br>position 2655 | CCATTTCAAGACGCTGAGGG |
| | R-primer – 8<br>position 3144 | AATGTTGCTCCACCAGTTGC |

| | | |
|---|---|---|
| | L-primer - 9 position 3066 | CTTGGGCGAGTCATACGAGA |
| | R-primer – 9 position 3599 | AGTCTGTTGGAAACGCTCAG |
| | L-primer – 10 position 3440 | AGCAGAGTCACCACGAACAT |
| | R-primer – 10 position 3895 | CGGTACTGATCAATCTTCTGCT |
| | L-primer – 11 position 3707 | TGATCCCAAGTAGTATCGTTCG |
| | R-primer – 11 position 3999 | ATGAGAGGAGGAGTCGATAGG |
| | L-primer – 12 Position 3895 | AGCAGAAGATTGATCAGTACCG |
| | R-primer – 12 Position 4397 | ATTCGCAAAGTGATGGAGGC |
| | L-primer – 13 position 4215 | TGGGATTTTAGCGAGAGGAGT |
| | R-primer – 13 position 4770 | AGATCACGTCCTAGTAGTGCT |
| | L-primer – 14 position 4582 | CACCTTCAGCTTGGCGTATA |
| | R-primer – 14 position 5031 | TCTTCCTCAGGTATGTCGCA |
| | L-primer – 15 position 4914 | TGTTGGCCCTTTTGAGTAGG |
| | R-primer – 15 position 5461 | TGTTGCTGCTGGTACTTCTG |
| | L-primer – 16 position 5257 | TCGTCATCAACATCAACAGGC |
| | R-primer – 16 position 5866 | TTTGAGGTCATATGGGCGGA |
| Thermocycling parameters (PCR): 1 cycle 94°C for 2 min, 30 cycles of 45 s at 94°C, 30 s at 56°C and 45 s at 72°C, and final extension for 2 min at 72°C – Taq DNA polymerase used | | |
| AfaDV<br><br>Oocyte and pyloric caeca detection<br><br>(PCR) | L-primer - 9 position 3066 | CTTGGGCGAGTCATACGAGA |
| | R-primer – 9 position 3599 | AGTCTGTTGGAAACGCTCAG |
| Thermocycling parameters (PCR): 1 cycle 98°C for 30 s, 35 cycles of 10 s at 98°C, 20 s at 66°C and 20 s at 72°C, and final extension for 2 min at 72°C - Q5 polymerase used | | |

| | | |
|---|---|---|
| AfaDV VP Cloning (PCR) | L-primer - position 252 restriction enzyme EcoRI | CGCgaattcATAGAAAAGGCTGTG |
| | R-primer position 3187 restriction enzyme HindIII | CGCaagcttCCTAATCCGCT |
| Thermocycling parameters (PCR): 1 cycle 98°C for 30 s, 30 cycles of 10 s at 98°C, 30 s at 66°C and 1 minute 30 s at 72°C , and final extension for 2 min at 72°C – Q5 polymerase used | | |
| SSaDV VP Cloning (PCR) | L-primer - position 2857 restriction enzyme HindIII | GGGGaagcttAGAAACCTAATCC |
| | R-primer position 5731 restriction enzyme EcoRI | CTGAgaattcCATTATGTCGGGTG |
| Thermocycling parameters (PCR): 1 cycle 98°C for 30 s, 30 cycles of 10 s at 98°C, 30 s at 67°C and 1 minute 30 s at 72°C, and final extension for 2 min at 72°C – Q5 polymerase used | | |
| AfaDV NS1, NS2, NS3 Cloning (PCR) | L-primer – 1 position 243 | TTTGAGGTCATATGGGCGGA |
| | R-primer – 8 position 3144 | AATGTTGCTCCACCAGTTGC |
| Thermocycling parameters (PCR): 1 cycle 98°C for 30 s, 30 cycles of 10 s at 98°C, 30 s at 67°C and 1 minute 30 s at 72°C, and final extension for 2 min at 72°C – Q5 polymerase used | | |
| SSaDV (PCR) | SSaDV_NS3_1_F | CAATACGCCGATTAGCTTACAG |
| | SSaDV_NS2_1_R_2 | TCCTCGCTCACTACTAATGTTG |
| Thermocycling parameters (PCR): 1 cycle 98°C for 30 s, 35 cycles of 10 s at 98°C, 30 s at 64°C and 40 s at 72°C , and final extension for 2 min at 72°C – Q5 polymerase used | | |
